# Transcriptomic stability or lability explains sensitivity to climate stressors in coralline algae

**DOI:** 10.1101/2021.04.18.440109

**Authors:** Tessa M. Page, Carmel McDougall, Ido Bar, Guillermo Diaz-Pulido

**Affiliations:** Griffth University School of Environment and Science and Australia Rivers Institute Nathan Campus, Griffith University, 170 Kessels Road, Nathan, QLD 4111, Australia; Environment Futures Research Institute Nathan Campus, Griffith University, 170 Kessels Road, Nathan, QLD 4111, Australia

## Abstract

Crustose coralline algae (CCA) are a group of calcifying red macroalgae crucial to tropical coral reefs because they form crusts that cement together the reef framework^1^. Previous research into the responses of CCA to ocean warming (OW) and ocean acidification (OA) have found reductions in calcification rates and survival^2,3^, with magnitude of effect being species-specific. Responses of CCA to OW and OA could be linked to evolutionary divergence time and/or their underlying molecular biology, the role of either being unknown in CCA. Here we show *Sporolithon durum*, a species from an earlier diverged lineage that exhibits low sensitivity to climate stressors, had little change in metabolic performance and did not significantly alter the expression of any genes when exposed to temperature and pH perturbations. In contrast, *Porolithon onkodes*, a species from a recently diverged lineage, reduced photosynthetic rates and had over 400 significantly differentially expressed genes in response to experimental treatments, with differential regulation of genes relating to physiological processes. We suggest earlier diverged CCA may be resistant to OW and OA conditions predicted for 2100, whereas taxa from more recently diverged lineages with demonstrated high sensitivity to climate stressors may have limited ability for acclimatisation.

## Main

Uncertainties associated with global change have presented challenges for predicting the persistence of species in the ocean. Transcriptomic profiling allows for investigation of molecular responses of organisms to stressors and can be informative in indicating mechanisms for resistance or adaptation^4^ such as tolerance^5-7^ and plasticity^6^, or can indicate sensitivity^5-7^. Resistance and adaptation as responses to climate stressors can be measured at a molecular level with transcriptomics and can be seen through transcriptomic plasticity (i.e., shifting expression profile of transcriptome) or as a muted or dampened transcriptomic response^6,8^. Phenotypic plasticity is a possible response to a changing environment, with the transcriptome being a phenotype that responds to environmental cues^6,9^, however, plasticity does not always indicate acclimatisation or adaptive strategy^10^. Environmental stressors can destabilise the transcriptome causing differential regulation of genes. This transcriptomic lability can be indicative of a deleterious stress outcome^6,11-13^. Conversely, a muted or dampened transcriptional response, which we refer to as transcriptomic stability, can indicate resistance^6,8,14^. Transcriptomic stability as a means for resistance to stressors has been documented in both gymnosperms (pines^11,12^) and angiosperms (tomato plants^13^ and *Arabidopsis thaliana*^5^). However, the prevalence of transcriptomic stability vs lability has not been investigated in arguably one of the most important group of “plants” in the ocean, crustose coralline algae (CCA). This may be particularly critical for understanding the molecular and cellular responses of marine algae to climate stressors.

CCA are important marine organisms because of their role as ecosystem engineers (e.g., construction of coral reefs and maerl beds^15^) and their contribution to the global carbon cycle^16^. Some CCA genera have persisted and diversified through times of elevated temperature and *p*CO_2_/reduced pH that equal or surpass levels projected for the year 2100^17-19^. One could speculate that the evolutionary environmental histories of some genera of CCA would result in inherent resistance to OW and OA, compared with recently diverged genera that have not been exposed to levels of seawater temperature and *p*CO_2_/pH that are more extreme than current ocean conditions^17,18^. Previous experiments have found CCA to be negatively impacted by OW and OA^2,20^, however, there is obvious variability in type and magnitude of response that seems to be species-specific. Whether this variability relates to divergent evolutionary histories remains unexplored. From previous literature, earlier diverging lineages tend to be more resistant to OW and OA^21,22^, whereas more recently diverged genera seem to be more sensitive^2,18,22,23^ (Fig. 1a,b, Supplementary Table 1). Investigating the molecular basis of these responses could reveal the mechanisms by which resistance is obtained and facilitate comparisons between species with differing evolutionary histories.

**Figure 1.**
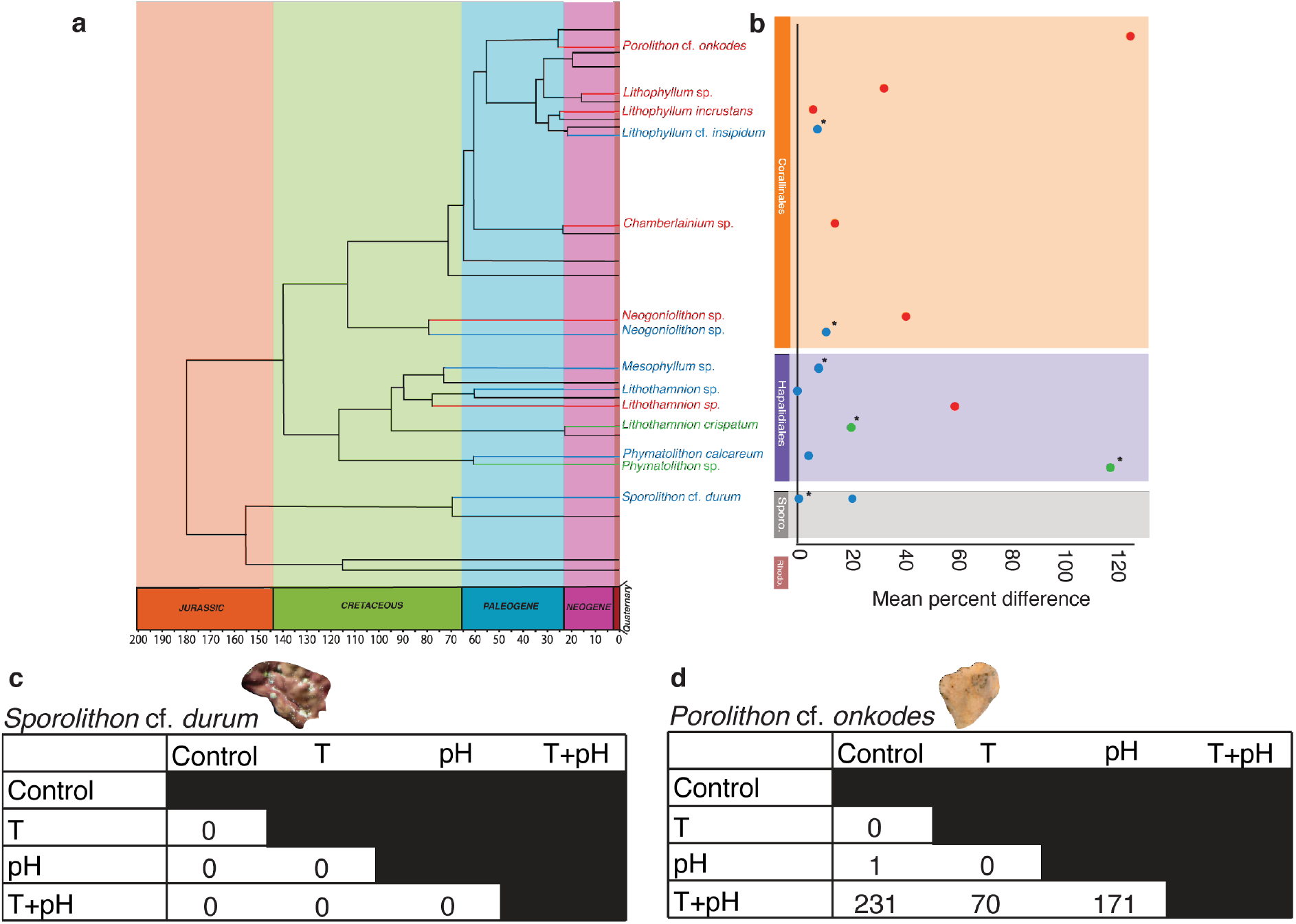
Variable responses of species from different orders of CCA to global change stressors from previous literature (a,b) and the current experiment (c,d). **a**, Phylogenetic tree, adapted from Peña, et al. ^18^, showing different species across orders of CCA and their estimated divergence time (x axis). Species names are colour coded to show direction of response to elevated temperature + reduced pH; red - significant negative response; green - significant positive response; blue - no significant effect. Data was obtained from 9 studies (Supplementary Table 1). **b**, Graphical representation of response of species from phylogenetic tree (Fig. 1a). Response is displayed as average relative change per species. Asterisks signify results from studies using Pulse-Amplitude-Modulation fluorometry to indicate photosynthesis, whereas all other studies directly measured dissolved O_2_ in seawater. from studies used in the reconstruction. **c**, Table of differentially expressed genes (DEGs) detected in pairwise comparisons from edgeR analyses across four treatments, control (27.2 °C and 8.0 pH), T (29.5 °C and 8.0 pH), pH (27.2 °C and 7.7 pH), and T+pH (29.5 °C and 7.7 pH), for *S*. cf. *durum* and **d**, *P*. cf. *onkodes*.

To do this, we exposed two divergent species, *Sporolithon* cf. *durum* and *Porolithon* cf. *onkodes*, to differing levels of temperature and pH, selected to reflect both current conditions and those projected for year 2100^19^. Experiments were conducted for 3 months. The following treatments were used: control (“control”: 27.2 °C and 8.0 pH), elevated temperature (“T”: 29.5 °C and 8.0 pH), reduced pH (“pH”: 27.2 °C and 7.7 pH), and combined stressors (“T+pH”: 29.5 °C and 7.7 pH). Physiological responses were measured, and RNA sequencing analysis was used to investigate transcriptomic stability or lability as a means to propose resistance or sensitivity in CCA. Transcriptomic response expression data was validated through RT-qPCR (see Methods).

The more recently diverged species, *P. onkodes*, showed transcriptomic lability, with 473 differentially expressed genes (DEGs) after 3 months in experimental treatments. *S. durum*, the earlier diverged lineage, had 0 DEGs, which we propose equates to transcriptomic stability. The transcriptional response in *P. onkodes* was only observed under the combined stressor treatment (T+pH; Fig. 1), only one gene (containing a lipoxygenase domain) was differentially expressed between the control and a single-stressor treatment (pH). The transcriptomic findings reflected physiological results, in which *S. durum* was proposed to be resistant to OW and OA in terms of survival and metabolic rates^21^, whereas net photosynthesis of *P*. cf. *onkodes* was significantly negatively affected under the combined treatment of T+pH (ANOVA, *F*_*1,16*_*= 4*.*782, p* = 0.046, Supplementary Fig. 1, Supplementary Table 2). The nature of the molecular response to OW and OA was unknown in CCA, but, as shown here, likely underlies potential resistance or susceptibility. Our study demonstrates that transcriptional response is also species-specific and the observed transcriptional stability or lability could be related to divergence time of CCA species.

The transcriptional response of *P. onkodes* likely reveals the molecular mechanisms underlying the observed physiological response to stress. 133 DEGs were uniquely found in the T+pH vs control comparison, and 27 were commonly differentially expressed across all treatments when compared to T+pH (Fig. 2a). Functional overrepresentation analysis of the 133 DEGs revealed biological processes relating to catabolism and metabolism of polysaccharides, plastid organisation, and phospholipid biosynthesis, with the latter two containing largest number of transcripts, 4 and 5 respectively. 17 transcripts were related to processes involving carbohydrates and lipids. Carbohydrates, specifically polysaccharides, have been suggested to play a role in the calcification process of CCA by acting as a matrix for calcification in their primary cell wall^24^. Changes in calcification rates observed in previous studies^2,23,25^ could potentially be explained by alterations of expression of these transcripts, with negative implications for reef formation and stability. Transcripts found across all T+pH comparisons were primarily overrepresented in biological functions relating to carbon acquisition and metabolism, suggesting the combination of elevated temperature and reduced pH results in changes to crucial, primary physical and chemical processes in *P. onkodes*.

**Figure 2.**
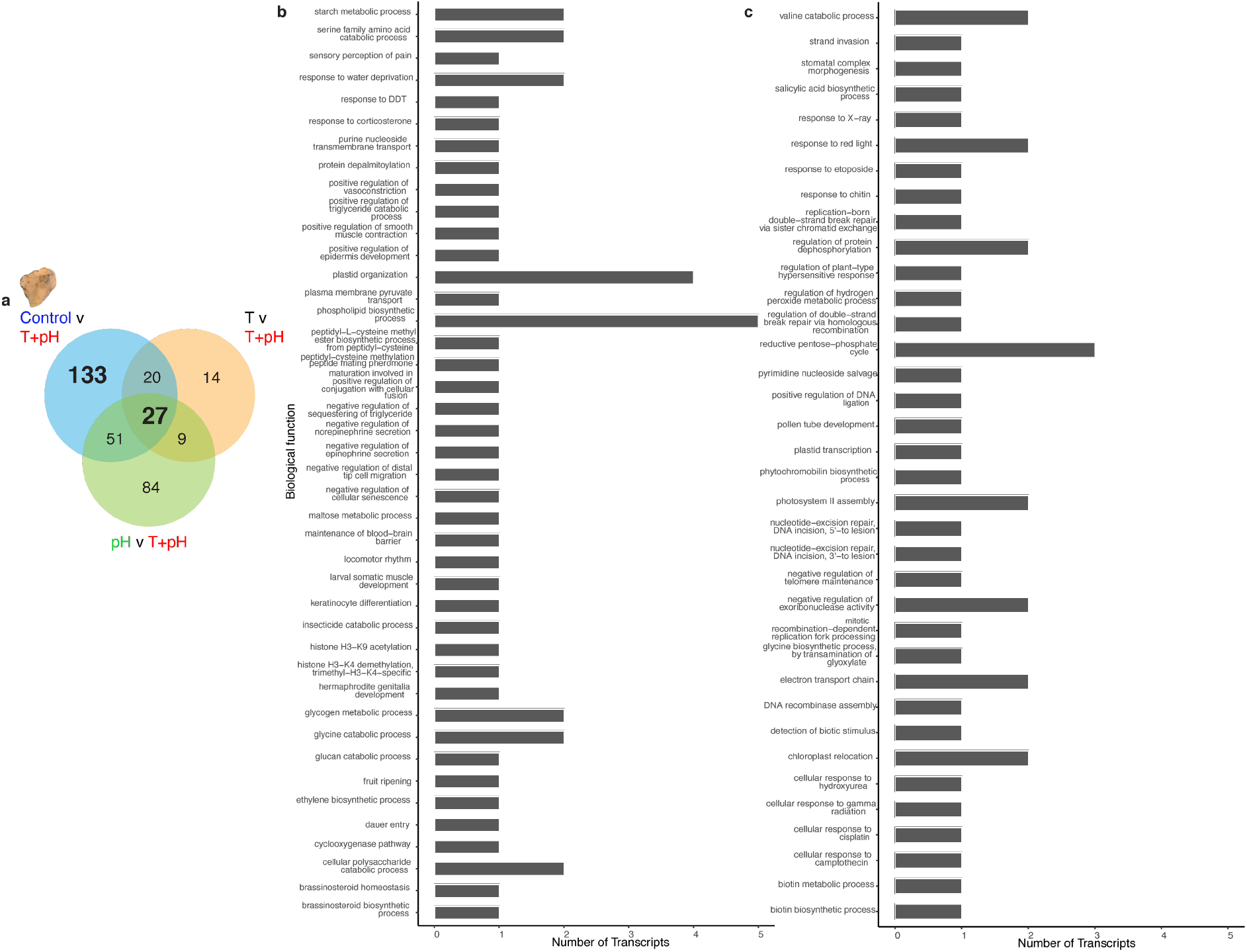
Unique and common differentially expressed genes found in pairwise comparisons between different experimental treatments. **a**, Venn diagram of DEGs found in pairwise comparisons for *P*. cf. *onkodes* for all treatments compared to the T+pH treatment. 27 common transcripts were found to be differentially expressed (DE) across all comparisons. **b**, Terminal nodes of overrepresented biological processes of the 133 transcripts that were found to be uniquely DE between the control treatment (27.2 °C and 8.0 pH) and the T+pH treatment (+2.3 °C and -0.3 pH units). Graph displays biological processes (y-axis) and number of transcripts per process (x-axis). **c**, Terminal nodes of overrepresented biological processes of the 27 shared transcripts that were found to be commonly DE across all comparisons. Graph displays biological processes (y-axis) and number of transcripts per process (x-axis).

Transcriptomic lability in *P. onkodes* was further observed in the patterns of differentially regulated genes within the T+pH treatment (Fig. 3a). Upregulated transcripts (130) were overrepresented in biological functional groups related to photorespiration, glycine metabolism, the reductive pentose-phosphate cycle, chloroplast organisation, and nucleotide-excision repair (Fig. 3b). Downregulated transcripts (99) were involved in biological functional groups related to the following mitochondrial processes: protein processing, positive regulation of membrane potential, stress-induced fusion, positive regulation of DNA replication, and calcium ion transport (Fig. 3c). Mitochondria are the powerhouses of eukaryotic cells and a growing area of plant research involves linking mitochondrial function and composition to environmental stress response^26,27^. Our finding of overrepresentation of mitochondrial-related processes in downregulated genes supports a role for mitochondria in the stress response and physiological processes of *P. onkodes*. Downregulation could be indicative of a negative effect on physiological processes, as was suggested in Antarctic algae in response to heat stress^28^.

**Figure 3.**
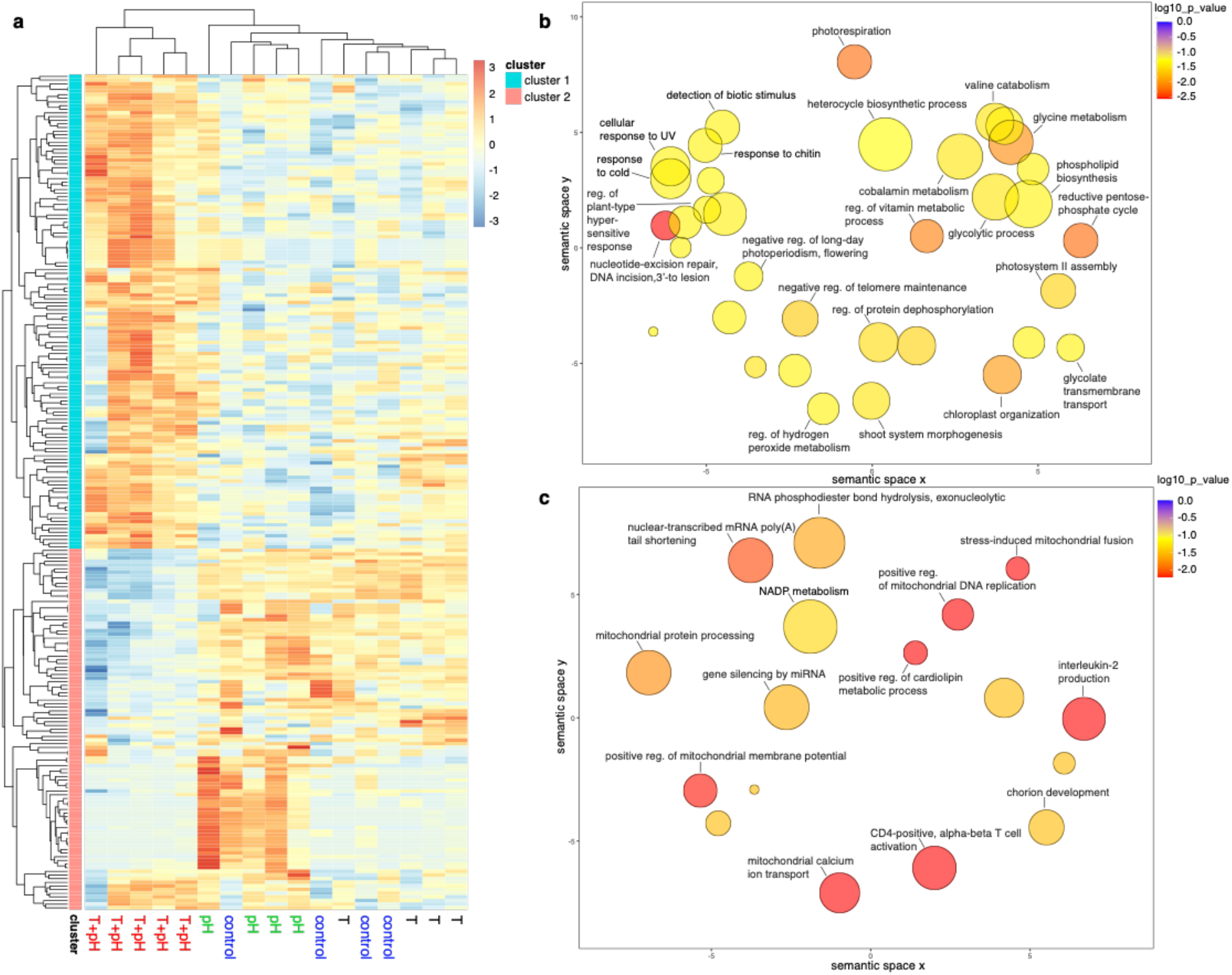
Patterns of differentially regulated gene expression across *P*. cf. *onkodes* experimental treatments. **a**, The T+pH treatment results in significantly upregulated and downregulated transcripts when compared to other treatments (heatmap of log2-fold-change FDR values < 0.05). Experimental treatments are labelled at the bottom of the heatmap. Two main clusters corresponding primarily to upregulated (cluster 1) and downregulated (cluster 2) transcripts in the T+pH treatment are evident. **b**, Overrepresented biological processes (terminal nodes; corrected p-values < 0.05) within cluster 1 transcripts include metabolism and catabolism, response to stimuli (biotic and abiotic), and regulation. Circle size indicates frequency of the gene ontology (GO) term in the UniProt database; colour indicates significance, on log^10^ *p* value scale. Axes have no intrinsic meaning; however, semantically similar GO terms remain closely together in the plot. **c**, Biological processes corresponding to terminal nodes from overrepresentation analysis of transcripts found in cluster 2 include mitochondrial processes, chorion development, and immune response.

A large proportion (51 of 130) of the transcripts that were found to be upregulated in the T+pH treatment encoded enzymes, and many of these are known components of the phosphate pentose pathway (PPP). This indicates that crucial metabolic processes of *P. onkodes* are affected by the synergistic effects of OW and OA. To visualise this, the proposed cellular locations of a subset of proteins encoded by transcripts from significantly enriched terminal biological processes from this study are visualised in Fig. 4 (all proteins listed in Supplementary Data File 2). Enzymes involved in the non-oxidative branch of the PPP were downregulated, whereas enzymes involved in the oxidative branch of the PPP were both downregulated (G6PDH and 6PGL) and upregulated (6PGDH). All differentially expressed enzymes involved in glycolytic reactions were upregulated (Fig. 4). Enzymes involved in Calvin-cycle specific reactions that were differentially expressed included PRK (significantly upregulated) and RuBisCO (downregulated, but not significantly). Two proposed thylakoid membrane proteins (cytochrome b_6_f and PGR5) were significantly upregulated; both of these proteins play a role in photosynthesis with involvement in either or both photosystem complexes. The mitochondrial proteins stomatin-2 and chaperone protein dnaJ were significantly downregulated; generally, upregulation of genes involved in protective stress responses (dnaJ) can facilitate a faster and more efficient response^8^. Collectively, these results indicate that global change drivers have a significant impact on the energy cycle of *P*. cf. *onkodes*. Enzymes involved in photorespiration were upregulated. Simultaneously, we found O_2_ production decreased. We propose that reallocation of energy to photorespiration resulted in a decrease in the efficiency of photosynthesis and this was observed in the decrease in the rate of net photosynthesis/average O_2_ production (Supplementary Figure 1).

**Figure 4.**
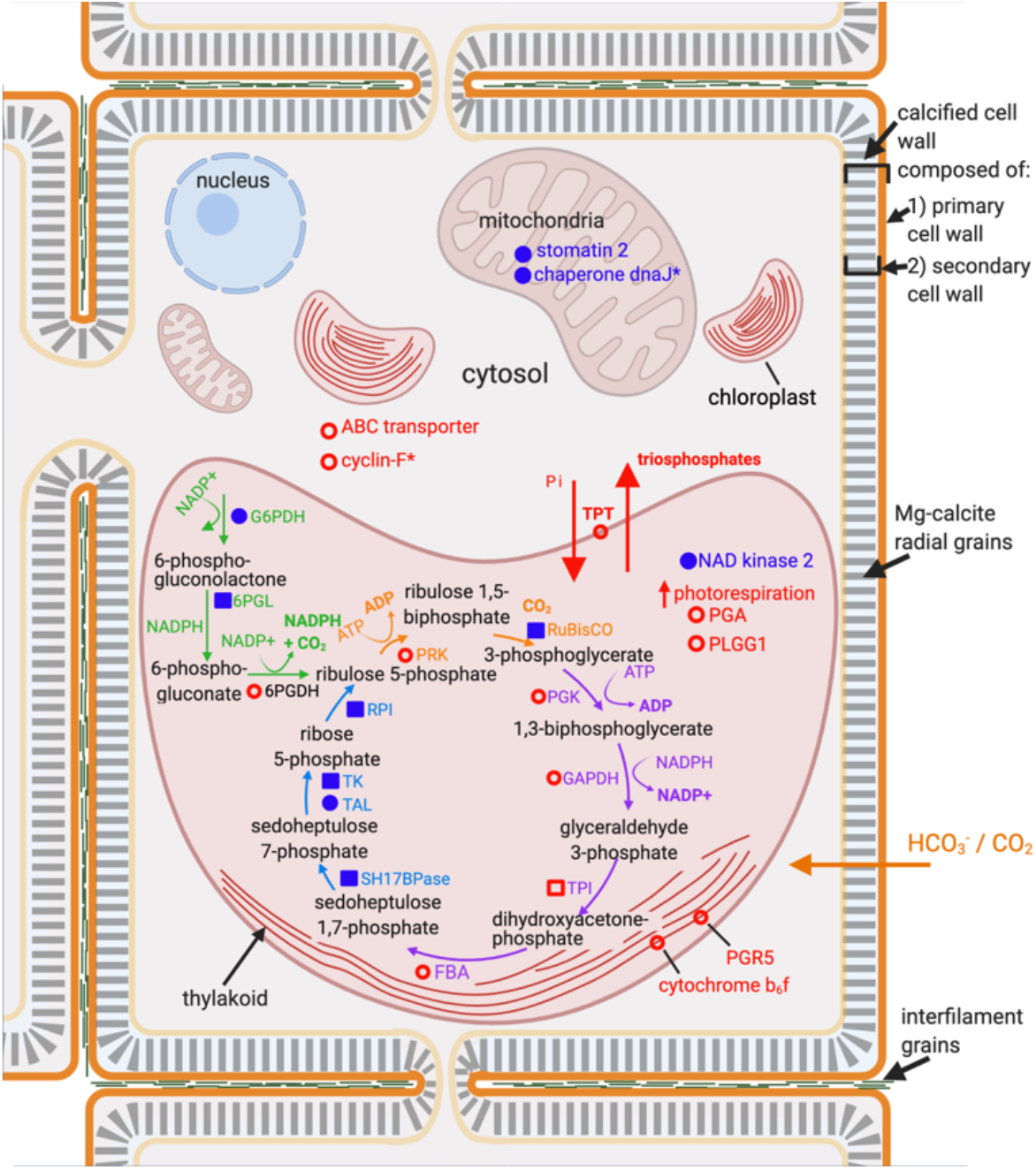
Conceptual model of the cellular pathways affected by global change stressors in *Porolithon* cf. *onkodes*. Conceptual model shows proposed subcellular locations of differentially expressed genes (DEGs) and proposed pathway involvement. Expression levels of proteins and enzymes are based on results from differential expression and functional overrepresentation analyses of transcripts within clusters 1 and 2 from Fig. 3a. Red open circles denote significant (FDR < 0.05) upregulation and blue solid circles significant downregulation under the T+pH treatment. Red open squares denote upregulated transcripts that were found within the *P. onkodes* transcriptome and within edgeR differential expression analysis but were not found to be significantly differentially expressed, and blue solid squares were downregulated, transcripts that were not significantly differentially expressed. * signify proteins that could have other subcellular localisations based on database (UniProt^29^ and COMPARTMENTS) investigations. Enzymes encoded by transcripts from this study were found in the Calvin-cycle (orange), glycolysis (purple), and the pentose phosphate pathway, both the oxidative (green) and nonoxidative branch (blue), with these pathways being proposed to occur within the plastid/chloroplast of the algae. Definitions of abbreviations for proteins are found in Supplementary Table 4. HCO_3_^-^ and CO_2_ are proposed to enter into the cell and directly used as a substrate for photosynthesis and calcification. Conceptual model was created with BioRender.com.

We propose that evolutionary history likely plays a role in resistance and susceptibility to stress amongst CCA species. To our knowledge, systematic studies that have specifically investigated differences in responses to stressors in the context of evolutionary history are lacking, however from the physiological studies available we suggest that divergence time could play a role in response (Fig. 1a,b). Studies from other taxa also support this theory. A review discussing temperature tolerance in terrestrial plants indicated a more tolerant species, *Arabidopsis thaliana*, had a muted transcriptional response compared with a less tolerant species, *Sorghum bicolor*^5^. Interestingly, these two species have divergent evolutionary histories, with the *Arabidopsis* genus having diverged ∼43 mya^30^ and the lineage containing *S. bicolor* estimated to have diverged between 3.9 – 2.4 mya^31^. Tolerance shown through transcriptomic stability or a muted transcriptomic response has been further documented in killifish that have evolved in high pollution conditions^7^. These findings are congruent with our hypothesis that evolutionary history may be a contributing factor to species-specific responses to stress. We propose that divergent evolutionary history may be an important driver behind the suggested resistance of *S. durum* to OA and OW, indicated in this study by transcriptomic stability and supported by prior studies^21^. On the other hand, the transcriptomic lability and negative effect of OW and OA on the physiology of *P. onkodes* suggests its overall sensitivity, which we propose is related to its more recent divergence. Future studies should continue to investigate the role of evolutionary history in response to climate stressors by systematically testing other possible contributing factors such as acclimatisation history and environmental history.

This study is the first to reveal differentially expressed genes and pathways that underpin physiological responses of CCA to stressors, and to implicate genes involved in crucial chemical and physical processes (i.e., PPP, glycolysis, Calvin-cycle, and photorespiration). We propose that the differing transcriptional responses of CCA to global change drivers provides an explanation into the species-specific responses of CCA observed in previous studies. Our study shows a polarised transcriptomic response to climate stressors in coralline algae grown under identical experimental conditions, which is further supported by differences in physiological response. We suggest transcriptomic plasticity or lability, as seen in *P. onkodes*, is indicative of susceptibility to global change drivers, whereas transcriptomic stability, as seen in *S. durum*, is indicative of resistance in CCA taxa. Although it may be argued plasticity is an expression of adaptation, that is not always the case^10^ and at times plasticity can be maladaptive or not contribute to increased resistance in an organism^6,10^. The findings from our study have implications for coral reefs worldwide. Our results indicate that *P*. cf. *onkodes*, an abundant reef-building species, may be negatively affected by predicted global change. In contrast, other tropical CCA species such as *S. durum*, although not currently major reef builders, may have the potential to thrive under predicted OW and OA scenarios.

## Methods

### Algae collection and experimental treatments

The two species used in this study were *Sporolithon* cf. *durum* and *Porolithon* cf. *onkodes*. These two species were chosen for the following reasons: 1) They are abundant, reef building species, with *P. onkodes* being the primary reef building species in the Great Barrier Reef (GBR), Australia; 2) *S. durum* and *P. onkodes* have been found to have different sensitives to global change drivers from previous studies^2,21^; 3) *S. durum* and *P. onkodes* come from genera with different evolutionary divergence times^18^; and 4) they are two of the only four CCA species that currently have sequenced transcriptomes^32^.

Adult fragments, ∼3 cm^2^, of CCA from the species *Sporolithon* cf. *durum* and *Porolithon* cf. *onkodes* were collected from lagoonal and reef crest sites surrounding Lizard Island, GBR, Australia. *S. durum* was collected between 7 – 9 m of depth and *P. onkodes* in depths no deeper than 3 m. Algal fragments were entirely crustose and were collected using hammer and chisel on SCUBA. Care was taken during collections to minimise impact on reef and distribute collections out over a wide area. After collection, algae fragments were transported to Lizard Island Research Station (LIRS) and held in an outdoor, flow through tank, maintaining similar seawater conditions to those measured at collection sites. Fragments were thoroughly cleaned of epiphytes within 24 hours of collection. Fragments that appeared to be very epiphytised (identified by large presence of burrowing worm holes and/or overabundance of epiphytic algae) were returned to collection sites. Fragments (*n* = 20 per species) were kept in control aquarium conditions for seven days at ambient temperature (26 °C), pH (8.00), salinity (35 ppt), and natural light (30 – 50 µmol quanta m^-2^ s-^1^ for *S. durum* and 140 µmol quanta m^-2^ s-^1^ for *P. onkodes*) prior to being placed into experimental treatments. Light was measured using an underwater quantum sensor LI-192 connected to a light meter LI-250 (LI-COR, USA).

Following the seven-days in common garden, fragments of CCA were exposed to 3 months of “control” (27.2 °C, 8.0 pH/450 µatm *p*CO_2_), “T” (29.5 °C, 8.0 pH/450 µatm *p*CO_2_), “pH” (27.2 °C, 7.7 pH/1000 µatm *p*CO_2_), or “T+pH” (29.5 °C, 7.7 pH/1000 µatm *p*CO_2_) treatments. Elevated temperature and *p*CO_2_ levels were selected to closely mimic future increases in temperature and *p*CO_2_ expected by the end of this century under the representative concentration pathway (RCP) 8.5^19^. In treatments where temperature and/or *p*CO_2_ were manipulated, levels were increased over 7 days reaching a 2.5 °C increase in temperature and 0.3 unit decrease in pH (∼ 1000 µatm *p*CO_2_). Temperature was maintained using titanium heaters (EcoPlus, Aqua Heat, 300 W), which were placed within each sump and were set to the desired temperature. Small 50 W glass heaters (Aqua One®) were placed in respective experimental tanks to correct for any heat loss when water moved from sump to experimental tanks. Temperature was adjusted daily across all treatments to mimic ambient temperature based on season (publicly available data from Australian Institute of Marine Science https://weather.aims.gov.au/#/station/1166) and +2.5 °C for elevated temperature treatments. pH/*p*CO_2_ was controlled by pH controllers (AquaController, Neptune Systems, USA) that injected either pure CO_2_ or ambient air into header sumps until set value was reached. Flow to each experimental tank was adjusted in the morning and evening, maintaining a flow rate of around 20 L hr^-1^, which allowed for complete turnover in experimental tanks twice an hour. Submersible water pumps (Aqua One® 8 W) were placed in each experimental tank to ensure adequate water flow. Mortality and changes in health were monitored and recorded throughout the experiment.

### Physiological measurements

Photosynthesis and respiration were measured for *P. onkodes* (pink morph) following techniques used in Page and Diaz-Pulido ^21^. Physiological data for *S. durum* was obtained from Page and Diaz-Pulido ^21^ in which similar treatments of elevated temperature and *p*CO_2_ were used however, conducted over a longer period of time (5 months versus 3 months in the current experiment). Physiological measurements of *P. onkodes* were normalised to surface area obtained through aluminium foil technique^33^.

### Seawater chemistry

Temperature and seawater pH (measured on total scale, pH_T_) were measured twice per day, at 08:00 and 15:00 in each experimental tank using a pH electrode with integrated temperature probe (Mettler Toledo, InLab Routine Pro) attached to a pH meter (Mettler Toledo, SevenGo Duo SG98). The pH electrode was calibrated on the total scale using Tris-HCl buffers^34^. Salinity was measured once a day using a conductivity meter (Mettler Toledo, SevenGo Pro). Total alkalinity (*A*_T_) was measured every 3 days for the first week of the experiment, and then every 6 – 7 days following using potentiometric titration on an automatic titrator (Mettler Toledo, T50) following standard operating procedures 3b^34^. pH_T_, *A*_T_, temperature, and salinity were used to calculate the remaining carbonate chemistry variables using the seacarb package version 3.2.12^35^ in the statistical software R version 3.5.1 (Supplementary Table 5). High-Mg calcite was calculated for a 16.4% MgCO_3_, following method described in Diaz-Pulido, et al. ^2^.

### Sampling and RNA extraction

Fragments of CCA were thoroughly cleaned prior to sampling for molecular analysis. Each fragment was examined under a microscope and all epiphytes, as far as possible, were removed. Directly before molecular sampling CCA were rinsed with filtered seawater and RNAlater® and then blotted with a kimwipe to remove bacterial film, following similar methods detailed in Page, et al. ^32^. Only the top, pigmented layer of the CCA fragments was collected under a microscope using sterile razors and placed directly into RNAlater®. Care was taken to avoid collection of any endolithics when obtaining samples. Samples were left at 4 °C for 24 hrs and then transferred to -20 °C prior to being transported to Griffith University for analysis. RNA extraction procedure followed the method detailed in Page, et al. ^32^. RNA quality and quantity were checked spectrophotometrically using an Invitrogen Qubit® Broad Range RNA kit. RNA yield ranged from 10.4 to 340 ng /µl. One RNA sample of *S. durum* yielded an undetectable amount of RNA and therefore was not used for further analyses. A random selection of samples, 2 – 3 samples from each treatment and species, were tested using the 4200 Tapestation System to ensure RNA quality and absence of degradation and contamination.

### cDNA synthesis and library preparation

All samples were diluted to reach a concentration of 25 ng/µl in 4.5 µl total volume prior to cDNA synthesis and library preparation. cDNA synthesis and library preparation followed single-cell sequencing (CEL-Seq2) protocols detailed in Hashimshony, et al. ^36,37^. Each sample (*n* = 39) was annealed to a specific primer that was designed with an anchored polyT, a unique barcode (6 bp), a unique molecular identifier (UMI) (increased to 7 bp), the 5’ Illumina adapter, and a T7 promoter. 1 µl of an external RNA control developed by the External RNA Controls Consortium (ERCC), Ambion® 4456740, at 1:10,000 dilution was added to each sample to control for variation in RNA expression that could be attributed to factors such as quality of starting material, platform or user error, and level of cellularity and RNA yield. mRNA was converted into DNA using SuperScript® II for cDNA synthesis and samples were pooled and then cleaned using AMPure XP® beads. Quality and concentration of RNA and DNA were checked throughout CEL-Seq2 protocol to ensure good quality and quantity both by Qubit® and Tapestation System. Final DNA concentration for the pooled library was 15.6 ng/µl.

### Sequencing

The pooled library was submitted to Ramaciotti Centre for Genomics, University of New South Wales, NSW, Australia. The library was cleaned up at Ramaciotti using AMPure XP® beads and passed additional quality control on both TapeStation and qPCR. Customised sequencing was performed on a single lane of an Illumina NovaSeq 6000, sequencing 26 bp on read 1, and 100 bp on read 2. A technical issue was identified for one sample for *P. onkodes*, and this sample was not considered in downstream analyses after sequencing.

### Sequence and differential expression analyses

Bioinformatic analyses were performed on Griffith University’s High Performance Computer Cluster, “Gowonda”, and statistical analyses and visualisations were performed using the statistical software R (v 3.6.1)^38^. Raw data were returned from Ramaciotti and processed using the Illumina BCL2FASTQ software, using default settings but with a minimum trimmed read length of 15. Quality control was performed on the FastQ files using FastQC (v 0.11.3, Babraham Bioinformatics). Reference transcriptomes were created using transcriptomes for *P. onkodes* and *S. durum*^32^. PolyA sequences were removed from reference transcriptomes using prinseq^39^. ERCC sequences, without polyA’s, were appended to the transcriptome files. Bowtie indices were generated using bowtie^40^ v 2 – 2.0.2. The resulting FASTA files were used to generate files mimicking gene transfer format (gtf) files following protocol laid out in McDougall, et al. ^41^. These “fake” gtf files were then used in the publicly available CEL-Seq-pipeline (https://github.com/yanailab/CEL-Seq-pipeline). Demultiplexed reads were mapped against the reference transcriptomes and read counts per transcript were generated. Samples that had less than 900,000 demultiplexed reads and had less than ∼ 40% mapped reads were removed from further analyses (*n* = 2 for *P. onkodes*) (Supplementary Table 6). Counts were imported into R (v 3.6.1) and corrected to account for the possibility of transcripts getting the same UMI. To correct for this and to convert UMI counts to transcript numbers the binomial method outlined in Grün, et al. ^42^ was used. Transcripts were filtered using a cut-off of 5 counts per transcript across all samples to remove transcripts with very low counts.

Differential gene expression analysis for each species was performed using the Bioconductor software package edgeR, v 3.16.8^43^. Negative binomial (NB) generalised linear models (GLMs) were fitted to transcript counts and common dispersions (0.169 and 0.293 for *P. onkodes* and *S. durum*, respectively) were estimated^43^. A design matrix of the experiment was used for analysis to identify expression in response to treatment. Quasi-likelihood (QL) F-tests were used in determining differential expression using default settings and the parameter ‘robust=TRUE’ to identify genes that were outliers from the mean-NB dispersion trend. Pairwise comparisons were conducted on specified constructed parameters (i.e. treatments) where genes that exhibited positive or negative log-fold changes were identified. Differentially expressed genes (DEGs) between treatments with a false discovery rate (FDR) cut-off of 5% and a log2-fold-change, looking at significantly expressed genes above a log2-fold-change of log_2_ (i.e., a fold-change of 1.2), were extracted and the datasets concatenated to use for downstream analyses and visualisations. FDR correction was applied using the Benjamini-Hochberg method on the *p*-values^43^. Count data were also analysed with DESeq2^44^ and pairwise comparisons were made to further validate edgeR analysis, DEG results were not significantly different between the two forms of analyses and *S. durum* still presented no significantly DEGs. To investigate DE expression further in *S. durum*, however, we used non-FDR corrected *p*-values to look at expression (Supplementary Figure 3).

Visualisation of DEGs was performed using variance stabilising transformed (vst) counts from edgeR. Principal component analyses were performed for each species to explore the variation in DEGs within species (Supplementary Figure 2). Heatmaps were constructed for DEGs from each species using the R package pheatmap (v 1.0.12)^45^. An FDR cut-off of 0.05 was used when creating the heatmap. Functional overrepresentation analysis of differentially expressed transcripts was performed in the Cytoscape^46^ plugin BiNGO^47^, where hypergeometric tests of gene ontology (GO) categories, specifically “biological process”, were used, with the annotated transcriptome of *P. onkodes* as a ‘background’, and a *p*-value (Benajmini-Hochberg FDR correction) cut-off of 0.01. BiNGO also allowed for identification of terminal node biological processes. REVIGO^48^ was used to summarise and visualise gene ontology terms obtained from enrichment analysis.

Similarity searches for DEGs were conducted using NCBI’s (National Center for Biotechnology Information) Basic Local Alignment Search Tool (BLAST)^49^ using the default e-value cut-off of 0.01. In order to identify KEGG pathway components we used the KEGG Mapper – Reconstruct Pathway tool^50^. KEGG annotations were obtained for all expressed and differentially expressed genes for each species using previous KEGG annotations from previously annotated transcriptomes^32^. Proposed cellular locations and pathway involvement of DEGs used in Fig. 4 were based on BLASTX similarity searches and KEGG Mapper Reconstruction results. Subcellular localisations were obtained through BLASTX and further checked on the subcellular localisation database, COMPARTMENTS (https://compartments.jensenlab.org/).

### RT-qPCR validation of expression results from CEL-Seq analysis

For qPCR validation of CEL-Seq edgeR expression analysis, reference genes and genes of interest (GOI) were chosen from edgeR normalised reads data. To obtain reference genes, we used a coefficient variation (CV) model to assess the degree of variation for each gene from edgeR normalised reads data^51^. To calculate the CV, we found the ratio of the standard deviation to the mean of each gene, high CVs indicate more variation in expression of a gene, whereas low CVs indicate low variation. Reference genes were chosen if CV < 0.47 and also had a low standard deviation, and GOIs were chosen by looking for the highest CVs. Additionally, we performed reciprocal BLAST analyses with the transcriptomes of *S. durum* and *P. onkodes* to find orthologues of reference genes that have been used with other species of red algae (i.e., the fleshy, red alga *Pyropia haitanensisi*), such as ß-tubulin and ubiquitin conjugating enzyme (UBC), and glyceraldehyde 3-phosphate dehydrogenase (GAPDH)^52^. GAPDH was only used as a reference gene for *S. durum* as it was found to have a low CV, however, GAPDH was used as a GOI for *P. onkodes* because it was found to be significantly differently expressed, with a high CV. Primers for GOIs were obtained from exploring *P. onkodes* edgeR data for highly, differentially expressed genes with large CVs, and then performing reciprocal blast in the *S. durum* transcriptome to find orthologues. Once genes were selected, primer sets were designed for reference genes and GOIs using Primer3^53^.

To test and optimise primer sets, cDNA was synthesised using Superscript III reverse transcriptase (Invitrogen**™)**) from DNase treated RNA of *S. durum* and *P. onkodes* and pooled for each species. Pooled cDNA was used as a template for PCR. PCRs using a thermal gradient (55 °C – 65 °C) were conducted to test primers and identify optimal annealing temperatures of primer sets. PCRs were run with 0.5 µL of forward and reverse primer (2.5 µM), 4.5 µL QuantiNova SYBR^®^ Green PCR Master Mix, 3.5 µL DNase/RNAse-Free H_2_O (PCR grade), and 1 µL cDNA pooled template (1:30 dilution). The thermal profile for PCR was 95 °C for 2 min, followed by 60 cycles of 95 °C for 5 s, thermal gradient (55 °C – 65 °C) for 30 s, and 60 °C for 10 s. PCRs were assessed by gel electrophoresis on a 2.5% agarose gels.

Subsequent RT-qPCR (CFX96 Touch™, Real-Time PCR Detection System, Bio-Rad) validation of reference genes and GOI was performed (Supplementary Table 7). Optimal dilution for each gene, primer efficiencies, and coefficient of determination (R^2^) were obtained from serial dilutions of standard curves for each primer set (Supplementary Table 7). The thermal profile for RT-qPCR was 95 °C for 2 min, followed by 60 cycles of 95 °C for 5 s, annealing temperature for specific primer set (identified from PCR) for 30 s, and 60 °C for 10 s, followed by a melt curve (65 °C to 95 °C in 0.5 °C increments for 5 s at each increment). RT-qPCR was then run for sample expression analysis. No template controls were included for each primer set and each sample and reactions were carried out in technical and biological triplicates. Suitable reference genes were calculated using geNorm^54^ based on geometric means to determine which reference candidates would be used as reference for expression analysis. Expression analysis was carried out in BioRad’s CFX Maestro Software, log_2_ Δ Δ Ct values (relative expression) for each sample were calculated and *p* values obtained to assess significant differential expression across treatments and individuals (Supplementary Figures 4 & 5), values were normalised to reference genes and predetermined efficiencies of each primer were entered based on standard curves (Supplementary Table 7).

### Systematic review of previous research and phylogenetic tree reconstruction

A systematic search for studies that investigated the metabolic responses of species of CCA to elevated temperature and ocean acidification (in combination) was conducted. The literature search was performed in the databases Google Scholar and Web of Science using keywords or topic codes such as ‘*crustose coralline algae*’ or ‘*coralline algae*’ in combination with ‘*photosynthesis, metabolic rates, ocean acidification, ocean warming, elevated temperature, reduced pH, elevated pCO*_*2*,_ *global change or climate change*’. We focused on studies that measured photosynthesis using a similar methodology as that used in the current study and in Page and Diaz-Pulido ^21^, however, as there were limited studies that fit our criteria, we supplemented the dataset with studies that used pulse amplitude modulated (PAM) fluorometry to determine photosynthetic capacity (Supplementary Table 1). The mean values of net photosynthesis or PAM fluorescence in the control and combined stressor treatments were obtained from publicly available datasets or, if results were only graphically represented, the built-in ruler and grids in Adobe Acrobat Pro DC *v* 2021.001.20138 (Adobe©) were used to obtain numerical values. Mean percent difference was calculated between control and the combined stressor treatment of elevated temperature and *p*CO_2_/reduced pH for each study (Supplementary Table 1). The absolute values of the percent differences were used in graphical representation. If studies manipulated other variables (i.e., nutrients or light), the control conditions for those variables were used. If studies measured over seasons, the average values from control and combined were taken across seasons. Mean percent differences were graphically represented adjacent to a reconstructed phylogenetic tree displaying species found in this review. The phylogenetic tree was adapted from Peña, et al.^18^.

### Statistical analyses for physiological data

Physiological data was analysed in R (v 3.6.1). Data were tested for normality through graphical analyses of residuals, using QQ normality plots. Data were log transformed if they did not meet normality. Two-way ANOVAs were run for photosynthesis and respiration data using temperature and pH as fixed factors. If a significant interaction between treatments was identified, ANOVAs were followed by Tukey’s HSD post hoc pairwise comparisons.

## Supplementary Information

Supplementary Information is available for this paper.

## Data availability

The datasets generated for this study can be found at https://osf.io/2nkr4/?view_only=08620a44c3534723b94e1c5c9bdd3bb0.

## Acknowledgements

This study was primarily funded by Australian Research Council grant DP160103071 awarded to G.D-P and in part by funds from the PADI Foundation awarded to T.M.P. The authors would like to acknowledge Griffith University’s High Performance Computer Cluster, “Gowonda”, and Indy Silva for assistance using Gowonda. We also acknowledge Ellie Bergstrom, Alexander Carlson, and Alea Laidlaw for their assistance during the experiment and for day-to-day maintenance. The authors also thank Dr. David Lambert and Dr. Sally Wasef for allowing us access to the TapeStation. We also thank the directors and maintenance staff of Lizard Island Research Station for their assistance and continued support throughout the duration of this study.

## References

1 Adey, W. H. Coral reefs: Algal structured and mediated ecosystems in shallow, turbulent, alkaline waters. J Phycol 34, 393–406, doi:10.1046/j.1529-8817.1998.340393.x (1998).

2 Diaz-Pulido, G., Anthony, K. R. N., Kline, D. I., Dove, S. & Hoegh-Guldberg, O. Interactions between ocean acidification and warming on the mortality and dissolution of coralline algae. J Phycol 1, 32–39, doi:10.1111/j.1529-8817.2011.01084.x (2012).

3 Cornwall, C. E., Diaz-Pulido, G. & Comeau, S. Impacts of ocean warming on coralline algal calcification: Meta-analysis, knowledge gaps, and key recommendations for future research. Front Mar Sci 6, 186, doi:10.3389/fmars.2019.00186 (2019).

4 Mohr, H. & Schopfer, P. in Plant Physiol 539–566 (Springer Berlin Heidelberg, 1995).

5 Raju, S. K. K., Barnes, A. C., Schnable, J. C. & Roston, R. L. Low-temperature tolerance in land plants: Are transcript and membrane responses conserved? Plant Sci 276, 73–86, doi:10.1016/j.plantsci.2018.08.002 (2018).

6 Debiasse, M. B. & Kelly, M. W. Plastic and evolved responses to global change: What can we learn from comparative transcriptomics? J Hered 107, 71–81, doi:10.1093/jhered/esv073 (2016).

7 Whitehead, A., Triant, D. A., Champlin, D. & Nacci, D. Comparative transcriptomics implicates mechanisms of evolved pollution tolerance in a killifish population. Mol Ecol 19, 5186–5203, doi:10.1111/j.1365-294X.2010.04829.x (2010).

8 Rivera, H. E. et al. A framework for understanding gene expression plasticity and its influence on stress tolerance. Mol Ecol, doi:10.1111/mec.15820 (2021).

9 Oostra, V., Saastamoinen, M., Zwaan, B. J. & Wheat, C. W. Strong phenotypic plasticity limits potential for evolutionary responses to climate change. Nat Commun 9, doi:10.1038/s41467-018-03384-9 (2018).

10 Grether, G. F. Environmental change, phenotypic plasticity, and genetic compensation. Am Nat 166, E115–E123, doi:10.1086/432023 (2005).

11 Gaspar et al. Comparative transcriptomic response of two Pinus species to infection with the pine wood nematode Bursaphelenchus xylophilus. Forests 11, 204, doi:10.3390/f11020204 (2020).

12 Fielding, N. & Evans, H. The pine wood nematode Bursaphelenchus xylophilus (Steiner and Buhrer) Nickle (= B. lignicolus Mamiya and Kiyohara): An assessment of the current position. J For 69, 35–46, doi:10.1093/forestry/69.1.35 (1996).

13 Bita, C. E. et al. Temperature stress differentially modulates transcription in meiotic anthers of heat-tolerant and heat-sensitive tomato plants. BMC Genomics 12, 384, doi:10.1186/1471-2164-12-384 (2011).

14 Bay, R. A. & Palumbi, S. R. Transcriptome predictors of coral survival and growth in a highly variable environment. Ecol Evol 7, 4794–4803, doi:10.1002/ece3.2685 (2017).

15 Nelson, W. A. Calcified macroalgae - critical to coastal ecosystems and vulnerable to change: A review. Mar Freshw Res 60, 787, doi:10.1071/mf08335 (2009).

16 Van Der Heijden, L. H. & Kamenos, N. A. Reviews and syntheses: Calculating the global contribution of coralline algae to total carbon burial. Biogeosciences 12, 6429–6441, doi:10.5194/bg-12-6429-2015 (2015).

17 Hönisch, B. et al. The geological record of ocean acidification. Science 335, 1058–1063, doi:10.1126/science.1208277 (2012).

18 Peña, V. et al. Radiation of the coralline red algae (Corallinophycidae, Rhodophyta) crown group as inferred from a multilocus time-calibrated phylogeny. Mol Phylogenet Evol 150, 106845, doi:10.1016/j.ympev.2020.106845 (2020).

19 IPCC. in Summary for Policymakers. In: IPCC Special Report on the Ocean and Cryosphere in a Changing Climate (eds Hans-O Pörtner et al.) (In press, 2019).

20 Kuffner, I. B., Andersson, A. J., Jokiel, P. L., Rodgers, K. u. S. & Mackenzie, F. T. Decreased abundance of crustose coralline algae due to ocean acidification. Nat Geosci 1, 114–117, doi:10.1038/ngeo100 (2007).

21 Page, T. M. & Diaz-Pulido, G. Plasticity of adult coralline algae to prolonged increased temperature and pCO2 exposure but reduced survival in their first generation. PLoS One 15, e0235125, doi:10.1371/journal.pone.0235125 (2020).

22 Bergstrom, E. et al. Inorganic carbon uptake strategies in coralline algae: Plasticity across evolutionary lineages under ocean acidification and warming. Mar Environ Res 161, 105107, doi:10.1016/j.marenvres.2020.105107 (2020).

23 Martin, S. & Gattuso, J.-P. Response of Mediterranean coralline algae to ocean acidification and elevated temperature. Glob Chang Biol 15, 2089–2100, doi:10.1111/j.1365-2486.2009.01874.x (2009).

24 Bilan, M. I. & Usov, A. I. Polysaccharides of calcareous algae and their effect on the calcification process. Russ J Bioorg Chem 27, 2–16, doi:10.1023/a:1009584516443 (2001).

25 Qui-Minet, Z. N. et al. Combined effects of global climate change and nutrient enrichment on the physiology of three temperate maerl species. Ecol Evol 9, 13787–13807, doi:10.1002/ece3.5802 (2019).

26 Jacoby, R. P. et al. Mitochondrial composition, function and stress response in plants. J Integr Plant Biol 54, 887–906, doi:10.1111/j.1744-7909.2012.01177.x (2012).

27 Liberatore, K. L., Dukowic-Schulze, S., Miller, M. E., Chen, C. & Kianian, S. F. The role of mitochondria in plant development and stress tolerance. Free Radical Biology and Medicine 100, 238–256, doi:10.1016/j.freeradbiomed.2016.03.033 (2016).

28 Hwang, Y.-s., Jung, G. & Jin, E. Transcriptome analysis of acclimatory responses to thermal stress in Antarctic algae. Biochem Biophys Res Commun 367, 635–641, doi:10.1016/j.bbrc.2007.12.176 (2008).

29 Apweiler, R. et al. UniProt: The universal protein knowledgebase. Nucleic Acids Res 32, D158–D169, doi:10.1093/nar/gkw1099 (2004).

30 Beilstein, M. A., Nagalingum, N. S., Clements, M. D., Manchester, S. R. & Mathews, S. Dated molecular phylogenies indicate a Miocene origin for Arabidopsis thaliana. PNAS 107, 18724, doi:10.1073/pnas.0909766107 (2010).

31 Liu, Q., Liu, H., Wen, J. & Peterson, P. M. Infrageneric phylogeny and temporal divergence of sorghum (Andropogoneae, Poaceae) based on low-copy nuclear and plastid sequences. PLoS One 9, e104933, doi:10.1371/journal.pone.0104933 (2014).

32 Page, T. M., McDougall, C. & Diaz-Pulido, G. De novo transcriptome assembly for four species of crustose coralline algae and analysis of unique orthologous genes. Sci Rep 9, 12611, doi:10.1038/s41598-019-48283-1 (2019).

33 Marsh Jr, J. A. Primary productivity of reef-building calcareous red algae. Ecology 51, 255–263, doi:10.2307/1933661 (1970).

34 Dickson, A. G., Sabine, C. L. & Christian, J. R. Guide to best practices for ocean CO2 measurements. (North Pacific Marine Science Organization, 2007).

35 seacarb: seawater carbonate chemistry with R v. 3.2.12 (2019).

36 Hashimshony, T., Wagner, F., Sher, N. & Yanai, I. CEL-Seq: Single-cell RNA-Seq by multiplexed linear amplification. Cell Rep 2, 666–673, doi:10.1016/j.celrep.2012.08.003 (2012).

37 Hashimshony, T. et al. CEL-Seq2: Sensitive highly-multiplexed single-cell RNA-Seq. Genome Biol 17, 77, doi:10.1186/s13059-016-0938-8 (2016).

38 R: A language and environment for statistical computing. R Foundation for Statistical Computing (Vienna, Austria, 2014).

39 Schmieder, R. & Edwards, R. Quality control and preprocessing of metagenomic datasets. Bioinformatics 27, 863–864, doi:10.1093/bioinformatics/btr026 (2011).

40 Langmead, B., Trapnell, C., Pop, M. & Salzberg, S. L. Ultrafast and memory-efficient alignment of short DNA sequences to the human genome. Genome Biol 10, R25, doi:10.1186/gb-2009-10-3-r25 (2009).

41 McDougall, C., Aguilera, F., Shokoohmand, A., Moase, P. & Degnan, B. M. Pearl sac gene expression profiles associated with pearl attributes in the silver-lip pearl oyster, Pinctada maxima. Front Genet 11, 597459–597459, doi:10.3389/fgene.2020.597459 (2021).

42 Grün, D., Kester, L. & van Oudenaarden, A. Validation of noise models for single-cell transcriptomics. Nat Methods 11, 637–640, doi:10.1038/nmeth.2930 (2014).

43 Robinson, M. D., McCarthy, D. J. & Smyth, G. K. edgeR: a Bioconductor package for differential expression analysis of digital gene expression data. Bioinformatics 26, 139–140, doi:10.1093/bioinformatics/btp616 (2010).

44 Love, M. I., Huber, W. & Anders, S. Moderated estimation of fold change and dispersion for RNA-seq data with DESeq2. Genome Biol 15, 550, doi:10.1186/s13059-014-0550-8 (2014).

45 pheatmap: Pretty heatmaps v. 1.0.12 (2015).

46 Shannon, P. et al. Cytoscape: A software environment for integrated models of biomolecular interaction networks. Genome Res 13, 2498–2504, doi:10.1101/gr.1239303 (2003).

47 Heymans, K., Kuiper, M. & Maere, S. BiNGO: a Cytoscape plugin to assess overrepresentation of Gene Ontology categories in Biological Networks. Bioinformatics 21, 3448–3449, doi:10.1093/bioinformatics/bti551 (2005).

48 Supek, F., Bošnjak, M., Škunca, N. & Šmuc, T. yREVIGO summarizes and visualizes long lists of gene ontology terms. PLoS One 6, e21800, doi:10.1371/journal.pone.0021800 (2011).

49 Altschul, S. F., Gish, W., Miller, W., Myers, E. W. & Lipman, D. J. Basic local alignment search tool. J Mol Biol 215, 403–410, doi:10.1101/pdb.top17 (1990).

50 Kanehisa, M., Sato, Y., Kawashima, M., Furumichi, M. & Tanabe, M. KEGG as a reference resource for gene and protein annotation. Nucleic Acids Res 44, D457–D462, doi:10.1093/nar/gkv1070 (2016).

51 Zeng, J. et al. Identification and analysis of house-keeping and tissue-specific genes based on RNA-seq data sets across 15 mouse tissues. Gene 576, 560–570, doi:10.1016/j.gene.2015.11.003 (2016).

52 Li, B., Chen, C., Xu, Y., Ji, D. & Xie, C. Validation of housekeeping genes as internal controls for studying the gene expression in Pyropia haitanensis (Bangiales, Rhodophyta) by quantitative real-time PCR. Acta Oceanologica Sinica 33, 152–159, doi:10.1007/s13131-014-0526-2 (2014).

53 Rozen, S. & Skaletsky, H. Primer3 on the WWW for general users and for biologist programmers. Methods Mol Biol 132, 365–386, doi:10.1385/1-59259-192-2:365 (2000).

54 Vandesompele, J. et al. Accurate normalization of real-time quantitative RT-PCR data by geometric averaging of multiple internal control genes. Genome Biol 3, 1–12, doi:10.1186/gb-2002-3-7-research0034 (2002).

