## Supplementary File 1 for "Transcriptomic stability or lability explains sensitivity to climate stressors in coralline algae"

**Supplementary Data File 1** – Transcriptomic stability or lability explains sensitivity to climate stressors in coralline algae

Tessa M. Page[1*], Carmel McDougall[1], Ido Bar[1,2], Guillermo Diaz-Pulido[1*]

[1]Griffth University School of Environment and Science and Australia Rivers Institute, Nathan Campus, Griffith University, Brisbane, Queensland, Australia
[2]Environment Futures Research Institute, Griffith University, Brisbane, Queensland, Australia

**Supplementary Table 1.** Metabolic responses of crustose coralline algae to the combined effects of elevated temperature and $p$CO$_2$/reduced pH, as reported in previous studies. Species that were used in the phylogenetic tree are bolded, all studies in this table were used in the response graph in Figure 1. Green = positive response to elevated temperature and $p$CO$_2$/reduced pH, red = negative response to elevated temperature and $p$CO$_2$/reduced pH, and blue = no response. Included within the table are species names, order, estimated first occurrence of genera (obtained from Peña, et al. [1]), levels of temperature and $p$CO$_2$/pH relative to current at time of divergence, species collection location, methodology of metabolic rate measurement, calculated mean value for control and elevated temperature and $p$CO$_2$/reduced pH treatments, and respective study. PAM = Pulse Amplitude Modulation Fluorometry. GBR = Great Barrier Reef, Australia.

| Species | Order | Divergence time *ca.* (mya) | Historic level of T°C + *p*CO$_2$/pH | Location | Methodology | Mean at control | Mean at high | Study |
|---|---|---|---|---|---|---|---|---|
| ***Porolithon* cf. *onkodes*** | *corallinales* | 22 | lower | GBR | Dissolved O$_2$ | 21 | -5 | Anthony, et al. [3] |
| *Porolithon* cf. *onokdes* | *corallinales* | 22 | lower | GBR | PAM | 513 | 506 | Bergstrom, et al. [4] |
| ***Chamberlainium* sp.** | *corallinales* | 39 | lower | Korea | Dissolved O$_2$ | 0.35 | 0.4 | Kim, et al. [5] |
| ***Lithophyllum cabiochae*** | *corallinales* | 27 | lower | France | Dissolved O$_2$ | 0.9 | 0.61 | Martin, et al. [6] |
| ***Lithophyllum incrustans*** | *corallinales* | 27 | lower | France | Dissolved O$_2$ | 1.78 | 1.68 | Qui-Minet, et al. [7] |
| ***Lithophyllum* cf. *insipidum*** | *corallinales* | 27 | lower | GBR | PAM | 441 | 464 | Bergstrom, et al. [4] |
| ***Neogoniolithon* sp.** | *corallinales* | 105 | higher | Mexico | Dissolved O$_2$ | 2.91 | 1.73 | Vásquez-Elizondo and Enríquez [8] |
| ***Neogoniolithon fosliei*** | *corallinales* | 105 | higher | GBR | PAM | 406 | 447 | Bergstrom, et al. [4] |
| ***Phymatolithon lusitanicum*** | *hapalidales* | 60 | lower | Portugal | Dissolved O$_2$ | 1.1 | 2.4 | Sordo, et al. [9] |
| ***Phymatolithon calcareum*** | *hapalidales* | 60 | lower | France | Dissolved O$_2$ | 1.7 | 1.8 | Qui-Minet, et al. [7] |
| ***Mesophyllum / Melyvonnnea sp.*** | *hapalidales* | 71 | higher | GBR | PAM | 472 | 495 | Bergstrom, et al. [4] |
| ***Lithothamnion* sp.** | *hapalidales* | 75 | higher | Mexico | Dissolved O$_2$ | 0.48 | 0.19 | Vásquez-Elizondo and Enríquez [8] |
| ***Lithothamnion corallioides*** | *hapalidales* | 75 | higher | France | Dissolved O$_2$ | 2.15 | 2.15 | Qui-Minet, et al. [7] |
| *Lithothamnion proliferum* | *hapalidales* | 75 | higher | GBR | PAM | 511 | 412 | Bergstrom, et al. [4] |
| ***Lithothamnion crispatum*** | *hapalidales* | 75 | higher | Brazil | PAM | 0.456 | 0.55 | Muñoz, et al. [10] |
| ***Sporolithon* cf. *durum*** | *sporolithales* | 137 | higher | GBR | Dissolved O$_2$ | 2.3 | 1.8 | Page and Diaz-Pulido [2] |
| *Sporolithon* cf. *durum* | *sporolithales* | 137 | higher | GBR | PAM | 514 | 513 | Bergstrom, et al. [4] |

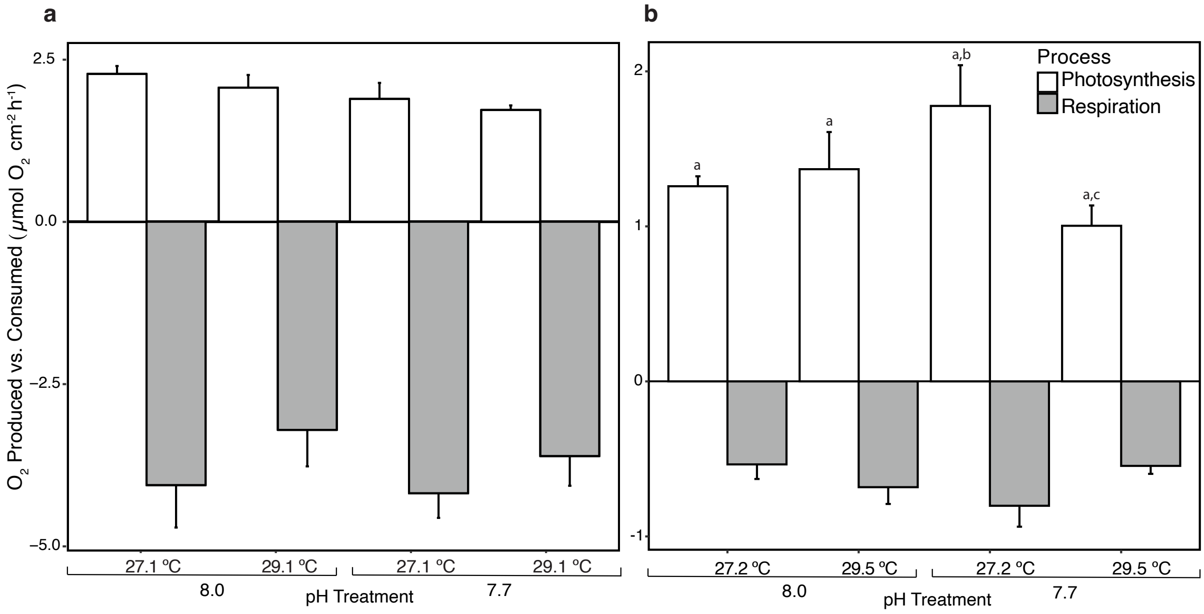


**Supplementary Figure 1.** Effect of global change drivers on metabolic processes of *Sporolithon* cf. *durum* and *Porolithon* cf. *onkodes*. **a**, Graph obtained from Page and Diaz-Pulido [2] of the effect of experimental treatment on the mean amount of $O_2$ produced or consumed by *S.* cf. *durum* after 5 months in treatment. Each bar represents mean $O_2$ produced or consumed ± standard error (SE), n = 5 -6. **b**, The effect of experimental treatment on the mean amount of $O_2$ produced and consumed by *P.* cf. *onkodes* after 3 months in treatment, values are means ± SE, n = 5. Significant differences ($p < 0.05$), resulting from Tukey HSD postdoc pairwise comparisons, are indicated by different lowercase letters.

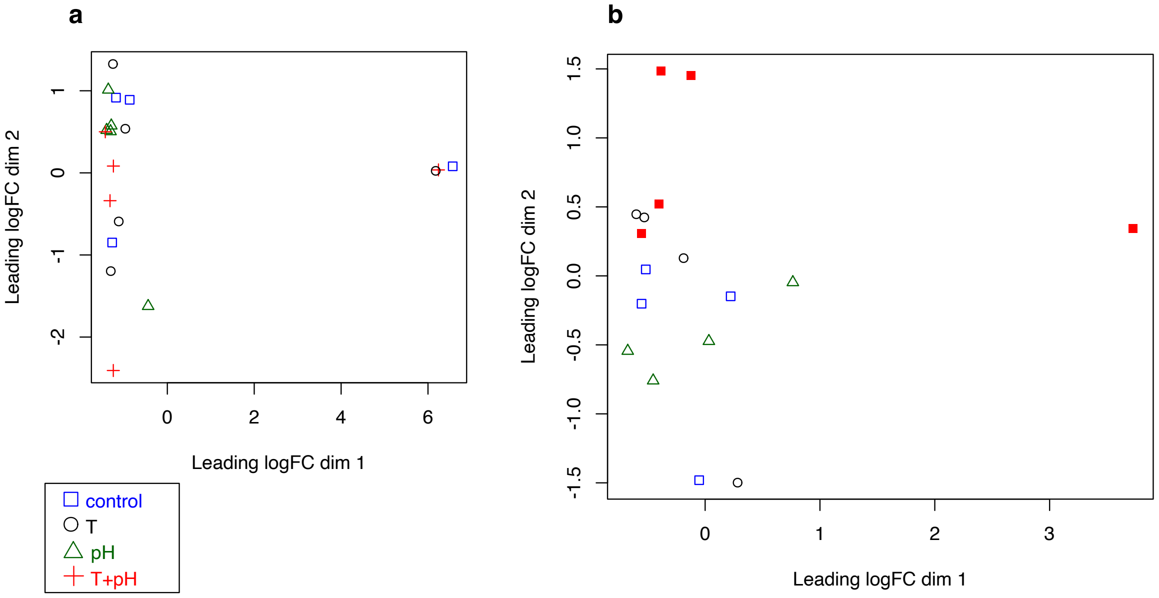


**Supplementary Figure 2.** Principal component analysis run on **a,** *Sporolithon* cf. *durum* and **b,** *Porolithon* cf. *onkodes*. Blue squares indicate the "control" treatment (8.0 pH and 27.2 ºC), black circles the "T" temperature (8.0 pH and 29.5 ºC), green triangles "pH" treatment (7.7 pH and 27.2 ºC), and the red plus sign the "T+pH" treatment  (7.7 pH and 29.5 ºC).

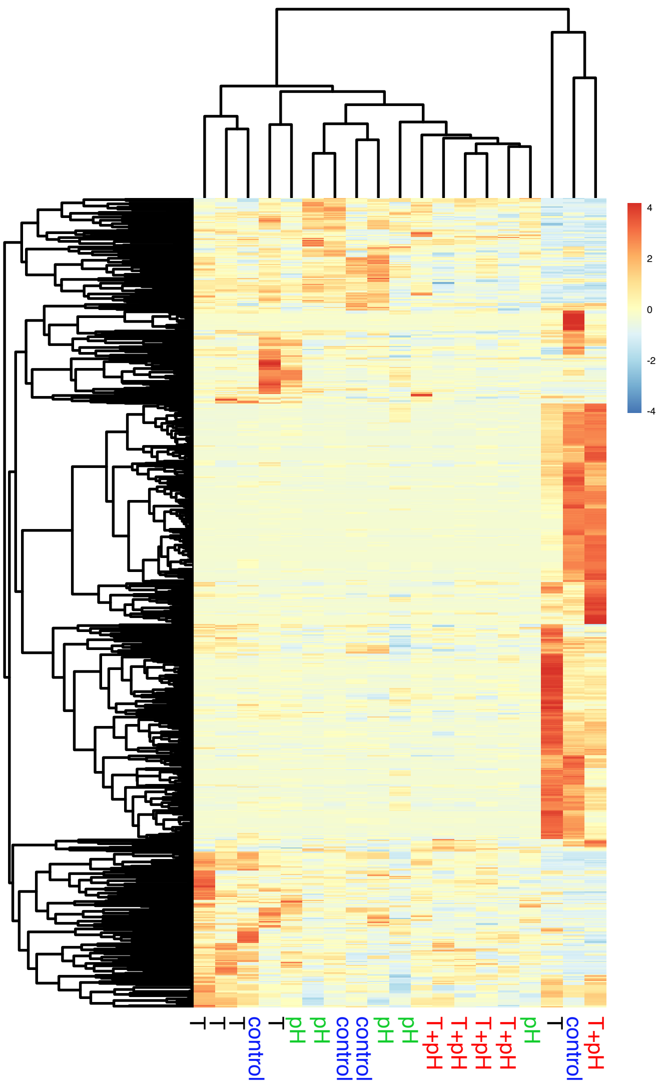


**Supplementary Figure 3.** A heatmap of differentially expressed genes from *Sporolithon* cf. *durum* based on *p* values < 0.05, without FDR adjustment. Heatmap shows expression of genes in *S.* cf. *durum*, however, these were not significant. Treatment labels correspond to the following treatments: control (27.2 ºC and 8.0 pH), T (29.5 ºC and 8.0 pH), pH (27.2 ºC and 7.7 pH), and T+pH (29.5 ºC and 7.7 pH).

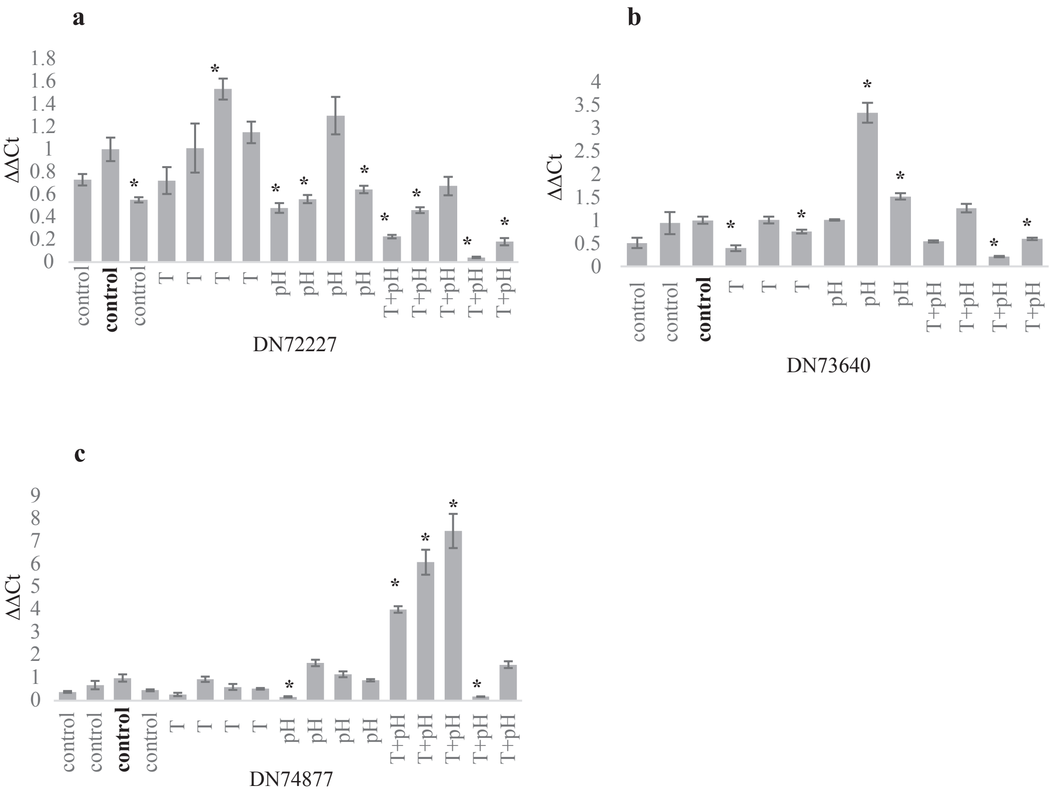


**Supplementary Figure 4.** RT-qPCR validation graphs for *Porolithon* cf. *onkodes*. * signify significant relative expression compared to a control sample (in bold). Values are the $log^2$ relative expression (ΔΔCt) of genes using two housekeeping genes as references (DN95780 and DN76782) ± normalised standard error of the mean (SEM). **a)** Transcript likely encoding for Hsp33 (BLASTX); **b)** transcript likely encoding photosystem II CP47 (BLASTX); and **c)** transcript likely encoding GAPDH (BLASTX).

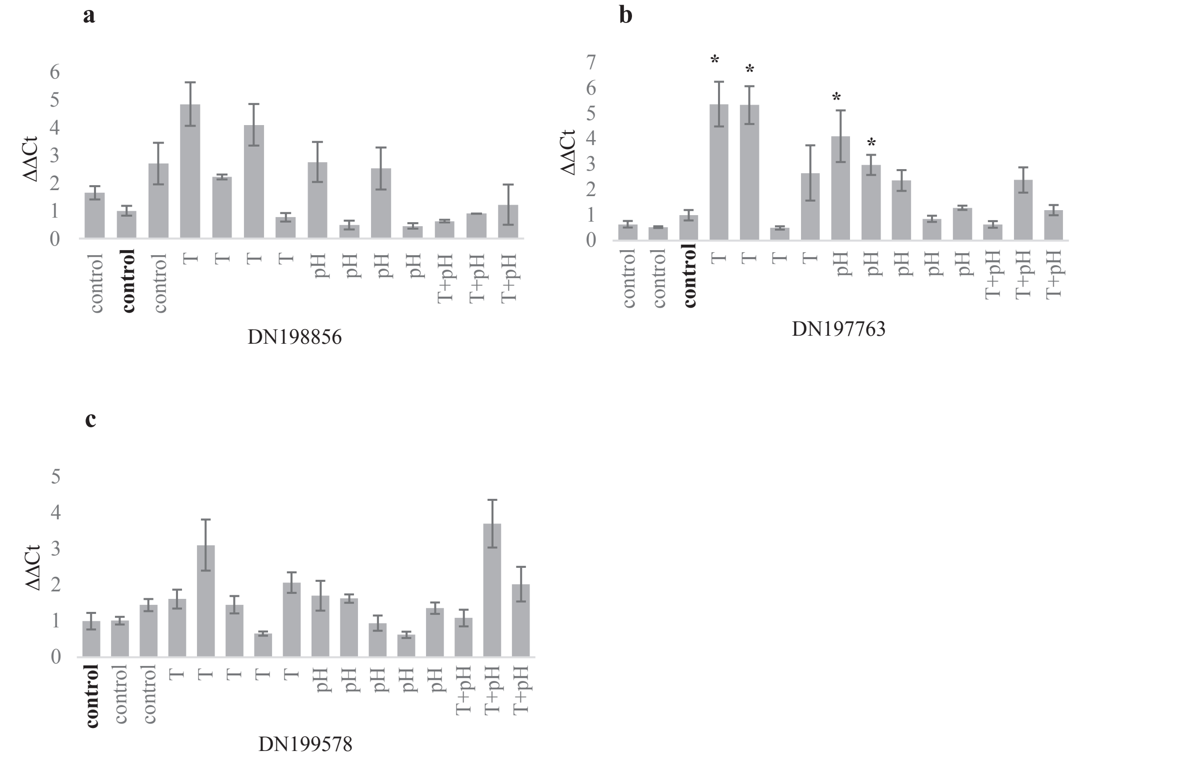


**Supplementary Figure 5.** RT-qPCR validation graphs for *Sporolithon cf. durum*. * signify significant expression compared to a control sample (in bold). Values are the log$^2$ relative expression (ΔΔCt) of genes using one reference (DN199936) ± normalised standard error of the mean (SEM). **a)** Transcript likely encoding for serine/threonine protein phosphate (BLASTX); **b)** transcript likely encoding transcript likely encoding Hsp33 (BLASTX); and **c)** transcript likely encoding acetyl-CoA (BLASTX).

**Supplementary Table 2.** Results of two-way analysis of variance (ANOVA) for the effects of temperature and pH ($p$CO$_2$) on the metabolic rates of *Porolithon* cf. *onkodes* (from the current experiment) and *Sporolithon* cf. *durum* (data acquired from Page & Diaz-Pulido, 2020).

| | | | Oxygen produced (µmol $O_2$ cm$^{-2}$ h$^{-1}$) | | | Oxygen consumed (µmol $O_2$ cm$^{-2}$ h$^{-1}$) | | |
|---|---|---|---|---|---|---|---|---|
| | Two-way ANOVA | Df | MS | *F* | *p* | MS | *F* | *p* |
| *Porolithon* cf. *onkodes* | Temperature | 1 | 0.073 | 4.443 | 0.054 | 0.015 | 0.294 | 0.595 |
| | pH | 1 | 0.000 | 0.000 | 0.988 | 0.021 | 0.405 | 0.534 |
| | Temperature * pH | 1 | 0.078 | 4.782 | **0.046** | 0.204 | 3.938 | 0.065 |
| | Residuals | 16 | 0.016 | | | 0.052 | | |
| *Sporolithon* cf. *durum* | Temperature | 1 | 0.056 | 0.439 | 0.516 | 3.026 | 1.830 | 0.192 |
| | pH | 1 | 1.217 | 9.560 | **0.006** | 0.495 | 0.299 | 0.591 |
| | Temperature * pH | 1 | 0.085 | 0.671 | 0.423 | 0.113 | 0.068 | 0.796 |
| | Residuals | 18 | 0.127 | | | 1.654 | | |

*Oxygen produced or consumed was normalised to surface area (cm$^2$) for *Porolithon* cf. *onkodes* and to the ash-free dry weight (g) of individual fragments for *S.* cf. *durum*.

**Supplementary Table 3.** Table of all proteins found to belong to terminal node biological processes from functional enrichment analysis. Table contains transcript/protein ID, top BLASTX similarity search result, proposed cell location based on BLASTX similarity search, expression pattern of transcript/protein based on ANOVA-like test in edgeR comparing all treatment combinations, cluster of heatmap (Fig. 3a), description of terminal node biological process obtained through enrichment analysis, and whether or not the transcript/protein was found to be unique to a specific biological process. All BLASTX hits were with red algae unless otherwise specified within table.

| Transcript /protein ID | BLASTX similarity result | Proposed cell location | Expression pattern | Heatmap cluster | Description of biological process | Unique to process |
|---|---|---|---|---|---|---|
| DN72649_c1_g2_i2 | phospho-ribulokinase | chloroplast | upregulated | 1 | detection of biotic stimulus, heterocycle biosynthetic process, reductive pentose-phosphate cycle, photosystem II assembly, chloroplast organisation, shoot system morphogenesis, regulation of hydrogen peroxide metabolism, regulation of protein dephosphorylation, regulation of plant-type hypersensitive response, response to chitin | N |
| DN74711_c1_g3_i1 | green algae hit, Trebouxia 2Fe-2S ferredoxin-like | | upregulated | 1 | heterocycle biosynthetic process, reductive pentose-phosphate cycle | N |
| DN70344_c0_g1_i2 | 6-phosphogluconate dehydro-genase NAD-binding domain-containing protein | cytosol | upregulated | 1 | valine catabolic process | Y |
| DN74502_c6_g1_i1 | serine-pyruvate amino-transferase | | upregulated | 1 | glycine metabolic process | Y |
| DN74877_c1_g1_i1 | glyceralde-hyde-3-phosphate dehydro-genase | chloroplast | upregulated | 1 | heterocycle biosynthetic process, reductive pentose-phosphate cycle, negative regulation of telomere maintenance, nucleotide-excision repair, DNA incision,3'-to lesion | N |
| DN76886_c0_g1_i1 | soybean hit, putative oxido-reductase | | upregulated | 1 | valine catabolic process | Y |
| DN70418_c0_g2_i2 | PsbB mRNA maturation | chloroplast | upregulated | 1 | heterocycle biosynthetic process, | N |

| | | | | | | |
|---|---|---|---|---|---|---|
| | factor Mbb1 | | | | photosystem II assembly, chloroplast assembly, shoot system morphogenesis, regulation of protein dephosphorylation | |
| DN75582_c0_g4_i1 | fructose-bisphosphate aldolase | | upregulated | 1 | glycolytic process | Y |
| DN75582_c0_g4_i2 | fructose-bisphosphate aldolase | | upregulated | 1 | glycolytic process | Y |
| DN74152_c0_g1_i4 | zinc finger protein 771, C2H2-type | | upregulated | 1 | heterocycle biosynthetic process | Y |
| DN71318_c0_g5_i1 | hyper-polarisation-activated voltage-gated potassium channel | | upregulated | 1 | regulation of vitamin metabolic process | Y |
| DN71318_c0_g5_i2 | hyper-polarisation-activated voltage-gated potassium channel | | upregulated | 1 | regulation of vitamin metabolic process | Y |
| DN74481_c13_g2_i1 | putative plastid 1-deoxy-D-xylulose 5-phosphate reducto-isomerase | | upregulated | 1 | phospholipid biosynthesis, response to cold | N |
| DN70500_c0_g1_i1 | transcription factor YY2 | | upregulated | 1 | heterocycle biosynthetic process | Y |
| DN74098_c0_g1_i1 | fructose-1,6-bisphosphate aldolase | chloroplast | upregulated | 1 | glycolytic process | Y |
| DN75667_c0_g1_i2 | phospho-glycerate kinase | chloroplast | upregulated | 1 | reductive pentose-phosphate cycle, glycolytic process | N |
| DN74365_c0_g1_i1 | pyridoxal kinase | | upregulated | 1 | phospholipid biosynthesis | Y |
| DN74365_c0_g1_i2 | pyridoxal kinase | | upregulated | 1 | phospholipid biosynthesis | Y |
| DN74961_c0_g1_i3 | glycerate translocator | chloroplast | upregulated | 1 | photorespiration, glycolate transmembrance transport, chloroplast assembly | N |
| DN71927_c0_g4_i1 | stem-loop binding protein of 41 kDa b | chloroplast | upregulated | 1 | chloroplast organisation, response to cold | N |
| DN67734_c4_g1_i1 | glycine dehydro-genase | mitochondrion | upregulated | 1 | glycine metabolic process | Y |
| DN71816_c0_g1_i2 | bacterial hit, GNAT fam | | upregulated | 1 | heterocycle biosynthetic process | Y |
| DN75856_c1_g1_i1 | triose-phosphate/ phosphate translocator | chloroplast | upregulated | 1 | cobalamin metabolism | Y |
| DN73854_c | cytochrome | chloroplast | upregulated | 1 | detection of biotic | N |

| | | | | | | |
|---|---|---|---|---|---|---|
| 3_g3_i2 | b6-f complex iron-sulfur subunit | | | | stimulus, photosystem II assembly, shoot system morphogenesis, regulation of hydrogen peroxide metabolism | |
| DN72323_c1_g3_i1 | serine hydroxy-methyl-transferase | mitochondrion or cytosolic | upregulated | 1 | photorespiration, glycine metabolic process, response to cold | N |
| DN70627_c0_g2_i2 | putative trans-criptional regulatory protein | | upregulated | 1 | heterocycle biosynthetic process | Y |
| DN70900_c0_g2_i1 | lysophospho-lipid acyl-transferase | | upregulated | 1 | phospholipid biosynthesis | Y |
| DN73638_c0_g1_i4 | ABC transporter B family member 5 | | upregulated | 1 | regulation of protein dephosphorylation | Y |
| DN73611_c0_g3_i1 | PGR5-like protein 1B | chloroplast | upregulated | 1 | heterocycle biosynthetic process | Y |
| DN76651_c3_g2_i3 | green algae hit, glyceralde-hyde-3-phosphate dehydro-genase | | upregulated | 1 | reductive pentose-phosphate cycle, glycolytic process, negative regulation of telomere maintenance, nucleotide-excision repair, DNA incision,3’-to lesion, cellular response to UV | N |
| DN76706_c0_g2_i2 | cob(I)yrinic acid a,c-diamide adenosyl-transferase | | upregulated | 1 | heterocycle biosynthetic process, cobalamin metabolism | N |
| DN75126_c0_g1_i3 | coral hit, 7,8 dihydro-8-oxoguanine tri-phosphatase-like | | upregulated | 1 | heterocycle biosynthetic process | Y |
| DN76392_c0_g1_i2 | probable phospho-lipase D | | upregulated | 1 | photorespiration | Y |
| DN72560_c0_g2_i1 | magnesium-chelatase subunit ChID | chloroplast | upregulated | 1 | heterocycle biosynthetic process | Y |
| DN77029_c1_g1_i1 | glycine de-carboxylase | mitochondrion | upregulated | 1 | glycine metabolic process | Y |
| DN75024_c0_g4_i1 | PsbP-like protein | chloroplast | upregulated | 1 | heterocycle biosynthetic process, negative regulation of long-day photoperiodism, flowering | N |
| DN74340_c0_g2_i1 | phospho-glycolate phosphatase 1B | chloroplast | upregulated | 1 | photorespiration | Y |
| DN72364_c | glycerol-3- | chloroplast | upregulated | 1 | phospholipid | Y |

| | | | | | | |
|---|---|---|---|---|---|---|
| 0_g1_i2 | phosphate acyl-transferase | | | | biosynthesis | |
| DN73854_c3_g3_i3 | phospho-ribulokinase | chloroplast | upregulated | 1 | detection of biotic stimulus, regulation of plant-type hypersensitive response, response to cold, response to chitin | N |
| DN76673_c2_g5_i1 | putative transporter | | upregulated | 1 | heterocycle biosynthetic process, cellular response to UV | N |
| DN74460_c0_g1_i1 | ATP phospho-ribosyl-transferase 2 | chloroplast | upregulated | 1 | heterocycle biosynthetic process | Y |
| DN76376_c1_g1_i3 | cyclin-F | | upregulated | 1 | heterocycle biosynthetic process, regulation of vitamin metabolic process | N |
| DN74914_c0_g2_i1 | unnamed | | downregulated | 2 | stress-induced mitochondrial fusion, mitochondrial calcium ion transmembrane transport, positive regulation of mitochondrial membrane potential, positive regulation of mitochondrial DNA replication, mitochondrial protein processing, positive regulation of cardiolipin metabolic process, interleukin-2 production, CD4-positive alpha-beta T cell activation | N |
| DN74914_c0_g2_i2 | stomatin prohibition-family | mitochondrion | downregulated | 2 | stress-induced mitochondrial fusion, mitochondrial calcium ion transmembrane transport, positive regulation of mitochondrial membrane potential, positive regulation of mitochondrial DNA replication, mitochondrial protein processing, positive regulation of cardiolipin metabolic process, interleukin-2 production, CD4-positive alpha-beta T cell activation | N |
| DN74914_c0_g2_i3 | stomatin prohibition-family | mitochondrion | downregulated | 2 | stress-induced mitochondrial fusion, mitochondrial calcium ion | N |

| | | | | | | |
|---|---|---|---|---|---|---|
| | | | | | transmembrane transport, positive regulation of mitochondrial membrane potential, positive regulation of mitochondrial DNA replication, mitochondrial protein processing, positive regulation of cardiolipin metabolic process, interleukin-2 production, CD4-positive alpha-beta T cell activation | |
| DN74986_c0_g1_i1 | NAD+ kinase | | downregulated | 2 | NADP metabolism | Y |
| DN71837_c1_g1_i2 | CCR4-NOT transcription complex | | downregulated | 2 | nuclear-transcribed mRNA poly(A) tail shortening, RNA phosphodiester bond hydrolysis, exonucleolytic, gene silencing by miRNA | N |
| DN71837_c1_g1_i9 | CCR4-NOT transcription complex | | downregulated | 2 | nuclear-transcribed mRNA poly(A) tail shortening, RNA phosphodiester bond hydrolysis, exonucleolytic, gene silencing by miRNA | N |
| DN76337_c0_g1_i4 | no hits | | downregulated | 2 | nuclear-transcribed mRNA poly(A) tail shortening, RNA phosphodiester bond hydrolysis, exonucleolytic, gene silencing by miRNA | N |
| DN74965_c0_g4_i1 | glucose-6-phosphate 1-dehydro-genase | chloroplast | downregulated | 2 | NADP metabolism | Y |
| DN76337_c1_g1_i1 | no hits | | downregulated | 2 | nuclear-transcribed mRNA poly(A) tail shortening, RNA phosphodiester bond hydrolysis, exonucleolytic, gene silencing by miRNA | N |
| DN69846_c0_g1_i5 | chaperone protein dnaJ | | downregulated | 2 | chorion development | Y |
| DN69846_c0_g1_i2 | chaperone protein dnaJ | | downregulated | 2 | chorion development | Y |
| DN76627_c1_g1_i1 | stomatin 2 | mitochondrion | downregulated | 2 | stress-induced mitochondrial fusion, mitochondrial calcium ion transmembrane, positive regulation of mitochondrial DNA replication transport, positive regulation of mitochondrial membrane potential, mitochondrial protein | N |

| | | | | | |
|---|---|---|---|---|---|
| | | | | processing, positive regulation of cardiolipin metabolic process | |
| DN69532_c0_g1_i2 | transaldolase | downregulated | 2 | NADP metabolism | Y |

**Supplementary Table 4.** Definition for abbreviations found in Fig. 4, conceptual model of *Porolithon* cf. *onkodes* cell.

| Abbreviation | Definition |
|---|---|
| PGA | Phosphoglycolate phosphatase |
| PGLGG1 | Plastidal glycolate/glycerate translocator |
| PPP | Pentose phosphate pathway |
| G6PDH | Glucose-6-phosphate 1-dehydrogenase |
| 6PGL | 6-phosphogluconolactonase |
| 6PGDH | 6-phosphogluconate dehydrogenase |
| RPI | Ribose 5-phosphate isomerase |
| TK | Transketolase |
| TAL | Transaldolase |
| SH17BPase | Sedoheptulose 1,7-biphosphatase |
| PRK | Phosphoribulokinase |
| RuBisCO | Ribulose-1,5-biphosphate carboxylase/oxygenase |
| PGK | Phosphoglycerate kinase |
| GAPDH | Glyceraldehyde-3 phosphate dehydrogenase |
| TPI | Triosephosphate isomerase |
| FBA | Fructose-bisphosphate aldolase |
| TPT | Triose phosphate/phosphate translocator |
| $P_i$ | Inorganic phosphate |
| PGR5 | Proton gradient regulation 5 |

**Supplementary Table 5.** Summary of mean carbonate chemistry in the experimental treatments. $pCO_2$, $HCO_{3-}$, and $CO_3^{2-}$ were calculated using the R package seacarb by inputting measured $pH_T$, total alkalinity (TA), temperature (Temp ºC), and a salinity of 35.5 ± 0.2. All values are mean ± standard error (SE). High-Mg calcite was calculated for 16.4% calcite following methods from Diaz-Pulido, et al. [11].

| Treatment [Target] | Temp ºC | $pH_T$ | TA µmol $kg^{-1}$ | $pCO_2$ µatm | $HCO_{3-}$ µmol $kg^{-1}$ | $CO_3^{2-}$ µmol $kg^{-1}$ | $\Omega_{High\text{-}Mg\ Calcite}$ |
|---|---|---|---|---|---|---|---|
| 27.2 ºC + pH 8.00 | 27.12 ± 0.060 | 8.00 ± 0.005 | 2291.49 ± 0.653 | 454.19 ± 8.178 | 1777.70 ± 4.330 | 209.14 ± 1.780 | 1.088 ± 0.010 |
| 29.5 ºC + pH 8.00 | 29.29 ± 0.071 | 7.99 ± 0.003 | 2290.71 ± 0.875 | 477.26 ± 3.982 | 1768.71 ± 2.980 | 212.72 ± 1.340 | 1.135 ± 0.030 |
| 27.2 ºC + pH 7.70 | 27.18 ± 0.051 | 7.70 ± 0.002 | 2290.91 ± 0.811 | 1020.44 ± 7.297 | 2005.62 ± 1.860 | 116.54 ± 7.970 | 0.606 ± 0.020 |
| 29.5 ºC + pH 7.70 | 29.48 ± 0.071 | 7.69 ± 0.003 | 2290.31 ± 0.722 | 1028.482 ± 7.076 | 1984.58 ± 1.540 | 125.02 ± 6.850 | 0.672 ± 0.030 |

**Supplementary Table 6.** Statistics for number of reads, counts, and mapping % generated during CEL-Seq pipeline for both *Sporolithon* cf. *durum* and *Porolithon* cf. *onkodes*.

| Species | Sample | # reads | # counts | % mapped |
|---|---|---|---|---|
| *S.* cf. *durum* | SD10_amb | 2792473 | 907968 | 67.25% |
| *S.* cf. *durum* | SD3_ph | 2905474 | 999343 | 69.65% |
| *S.* cf. *durum* | SD4_amb | 7230595 | 2078440 | 74.69% |
| *S.* cf. *durum* | SD5_ph | 17392054 | 5225552 | 72.16% |
| *S.* cf. *durum* | SD6_tph | 7571206 | 2565129 | 69.88% |
| *S.* cf. *durum* | SD7_tph | 3042944 | 948133 | 69.44% |
| *S.* cf. *durum* | SD8_temp | 3639126 | 1403593 | 53.51% |
| *S.* cf. *durum* | SD9_ph | 873800 | 262942 | 59.51% |
| *S.* cf. *durum* | SD1_temp | 4455442 | 1745783 | 67.34% |
| *S.* cf. *durum* | SD2_temp | 25755969 | 9499485 | 70.81% |
| *S.* cf. *durum* | SD11_ph | 9239774 | 3644235 | 69.84% |
| *S.* cf. *durum* | SD12_tph | 2052150 | 775382 | 65.55% |
| *S.* cf. *durum* | SD13_amb | 4700800 | 1497574 | 62.54% |
| *S.* cf. *durum* | SD14_ph | 13025010 | 2709167 | 73.76% |
| *S.* cf. *durum* | SD15_tph | 10015974 | 3138109 | 61.05% |
| *S.* cf. *durum* | SD17_amb | 12483294 | 3784547 | 71.55% |
| *S.* cf. *durum* | SD18_temp | 75371988 | 21542260 | 63.91% |
| *S.* cf. *durum* | SD19_tph | 14555346 | 2556820 | 71.85% |
| *S.* cf. *durum* | SD20_temp | 6520222 | 2365847 | 50.66% |
| *P.* cf. *onkodes* | PO10_amb | 11693266 | 2629626 | 49.28% |
| *P.* cf. *onkodes* | PO11_ph | 958480 | 159937 | 47.18% |
| *P.* cf. *onkodes* | PO12_tph | 1813570 | 1256101 | 51.77% |
| *P.* cf. *onkodes* | PO14_ph | 970230 | 177249 | 43.78% |
| *P.* cf. *onkodes* | PO15_tph | 16810993 | 3012292 | 57.63% |
| *P.* cf. *onkodes* | PO16_amb | 2638715 | 532955 | 39.93% |
| *P.* cf. *onkodes* | PO17_amb | 1338205 | 257316 | 51.17% |
| *P.* cf. *onkodes* | PO18_temp | 910868 | 191000 | 50.01% |
| *P.* cf. *onkodes* | PO1_temp | 1338887 | 244611 | 49.36% |
| *P.* cf. *onkodes* | PO2_temp | 13190073 | 2874863 | 50.56% |
| *P.* cf. *onkodes* | PO3_ph | 3767250 | 625636 | 57.24% |
| *P.* cf. *onkodes* | PO4_amb | 1209823 | 205975 | 56.76% |
| *P.* cf. *onkodes* | PO5_ph | 5228592 | 988016 | 49.91% |
| *P.* cf. *onkodes* | PO6_tph | 7691073 | 1530097 | 50.63% |
| *P.* cf. *onkodes* | PO7_tph | 5316967 | 861006 | 60.67% |
| *P.* cf. *onkodes* | PO8_temp | 4164386 | 739996 | 52.81% |

**Supplementary Table 7.** Transcripts used for RT-qPCR validation of CEL-Seq expression profiles. Primers for housekeeping (HK) genes and genes of interest (GOI) were designed and used for validation. Table has accession number for transcripts, species, BLASTX annotation of sequence, primer sequences, expected amplicon size (bp), primer optimal annealing temperature (Tm) ºC, and PCR efficiency (%) and coefficient of determination ($R^2$) from qPCR standard curve analysis. HK genes genes were chosen from commonly used HKGs, such as beta-tubulin (β-tubulin), glyceraldehyde 3-phosphate dehydrogenase (GAPDH) and ubiquitin C (UBC), and through investigation of transcripts with low standard deviation and that were not significantly differentially expressed through edgeR analysis (i.e., heme oxygenase).

| Trinity accession number | Species | BLASTX Annotation | Primer sequence (5'-3')[a] | Expected amplicon size (bp) | Tm (°C) | PCR efficiency (%) ($R^2$) |
|---|---|---|---|---|---|---|
| DN76782_c6_g1_i1 | *Porolithon* cf. *onkodes* | β-tubulin (HKG) | (F) TCGGCCCTACTGAGTCGATT<br>(R) CTGGAGAAGGCATGGACGAG | 182 | 57.2 | 99.8 (0.989) |
| DN95780_c0_g1_i1 | *Porolithon* cf. *onkodes* | heme oxygenase (HKG) | (F) AACCAGAATTACTTGTGTCGCA<br>(R) CTGTACCTTCATTGCCAGAAAGT | 117 | 55.1 | 96 (0.89) |
| DN74877_c1_g1_i1 | *Porolithon* cf. *onkodes* | GAPDH (GOI) | (F) TGTCATTGCTGGCGAGGATT<br>(R) CTTCGCTCCCGCCTGAATAT | 175 | 59.7 | 95.5 (0.91) |
| DN72227_c0_g4_i1 | *Porolithon* cf. *onkodes* | HSP33 (GOI) | (F) AGGTACGAACTTTGCGGTGT<br>(R) TGCCAAACCCATGCATTTCG | 166 | 57.2 | 100.1 (0.96) |
| DN73640_c0_g1_i1 | *Porolithon* cf. *onkodes* | photosystem II CP47 (GOI) | (F) CCTATGGACAAAGGCGATGG<br>(R) TGACGCTAACACCAACTTGC | 214 | 55.1 | 95 (0.99) |
| DN71939_c0_g6_i1 | *Porolithon* cf. *onkodes* | acetyl-CoA (GOI) | (F) ACTCCAACTTCAAACGTGCG<br>(R) ATCCACATTCACAGCACCGT | 224 | 59.7 | 120.5 (0.87) |
| DN73238_c10_g1_i1 | *Porolithon* cf. *onkodes* | ferritin-3, chloroplastic (GOI) | (F) ATCCACCTTCACAAGCCGAC<br>(R) AACGAAAACGAGCCCTGACA | 154 | 64 | 85.4 (0.99) |
| DN199936_c0_g1_i2 | *Sporolithon* cf. *durum* | GAPDH (HKG) | (F) ACAACTCGCAGGAAAGCTCA<br>(R) TGCTTTCATCGGTGCCCTTA | 212 | 55.1 | 120 (0.99) |
| DN198856_c0_g2_i4 | *Sporolithon* cf. *durum* | serine/threonine protein phosphatase (GOI) | (F) ATGCAGGCCCTTGAGTTTGT<br>(R) TGATACCGTTGCTCTGCCAG | 241 | 64 | 109 (-0.90) |
| DN199578_c0_g1_i2 | *Sporolithon* cf. *durum* | acetyl-CoA (GOI) | (F) AGACGGTGCAGTTGGAGATC<br>(R) ATATCCCCACCTTCCGATGC | 219 | 59.7 | 109 (0.96) |
| DN196675_c0_g3_i7 | *Sporolithon* cf. *durum* | UBC (HKG) | (F) TGCACAAACTACCTCACGCA<br>(R) ATGGTGCTCACTTGCTCACA | 192 | 57.2 | 88.9 (-0.94) |
| DN197763_c0_g2_i5 | *Sporolithon* cf. *durum* | HSP33 (GOI) | (F) CATCGGTCCAGGTCACTACG<br>(R) ATCGCGGCATAGAACTGAGG | 223 | 57.2 | 112.4 (-0.94) |

[a] F, forward primer; R, reverse primer.

### References


1 Peña, V. *et al.* Radiation of the coralline red algae (Corallinophycidae, Rhodophyta) crown group as inferred from a multilocus time-calibrated phylogeny. *Mol Phylogenet Evol* **150**, 106845, doi:10.1016/j.ympev.2020.106845 (2020).

2 Page, T. M. & Diaz-Pulido, G. Plasticity of adult coralline algae to prolonged increased temperature and *p*CO2 exposure but reduced survival in their first generation. *PLoS One* **15**, e0235125, doi:10.1371/journal.pone.0235125 (2020).

3 Anthony, K. R. N., Kline, D. I., Diaz-Pulido, G., Dove, S. & Hoegh-Guldberg, O. Ocean acidification causes bleaching and productivity loss in coral reef builders. *PNAS* **105**, 17442, doi:10.1073/pnas.0804478105 (2008).

4 Bergstrom, E. *et al.* Inorganic carbon uptake strategies in coralline algae: Plasticity across evolutionary lineages under ocean acidification and warming. *Mar Environ Res* **161**, 105107, doi:10.1016/j.marenvres.2020.105107 (2020).

5 Kim, J.-H. *et al.* Global warming offsets the ecophysiological stress of ocean acidification on temperate crustose coralline algae. *Mar Pollut Bull* **157**, 111324, doi:10.1016/j.marpolbul.2020.111324 (2020).

6 Martin, S., Cohu, S., Vignot, C., Zimmerman, G. & Gattuso, J.-P. One-year experiment on the physiological response of the Mediterranean crustose coralline alga, *Lithophyllum cabiochae*, to elevated $pCO_2$ and temperature. *Ecol Evol* **3**, 676-693, doi:10.1002/ece3.475 (2013).

7 Qui-Minet, Z. N. *et al.* Combined effects of global climate change and nutrient enrichment on the physiology of three temperate maerl species. *Ecol Evol* **9**, 13787-13807, doi:10.1002/ece3.5802 (2019).

8 Vásquez-Elizondo, R. M. & Enríquez, S. Coralline algal physiology is more adversely affected by elevated temperature than reduced pH. *Sci Rep* **6**, 19030, doi:10.1038/srep19030 (2016).

9 Sordo, L., Santos, R., Barrote, I. & Silva, J. Temperature amplifies the effect of high $CO_2$ on the photosynthesis, respiration, and calcification of the coralline algae Phymatolithon lusitanicum. *Ecol Evol* **9**, 11000-11009, doi:10.1002/ece3.5560 (2019).

10 Muñoz, P. T. *et al.* Short-term interactive effects of increased temperatures and acidification on the calcifying macroalgae *Lithothamnion crispatum* and *Sonderophycus capensis*. *Aquat Bot* **148**, 46-52, doi:10.1016/j.aquabot.2018.04.008 (2018).

11 Diaz-Pulido, G., Anthony, K. R. N., Kline, D. I., Dove, S. & Hoegh-Guldberg, O. Interactions between ocean acidification and warming on the mortality and dissolution of coralline algae. *J Phycol* **1**, 32-39, doi:10.1111/j.1529-8817.2011.01084.x (2012).
